# The AAA+ ATPase RUVBL2 is a dual switch controlling invadosome formation and architecture

**DOI:** 10.64898/2026.11.20.768112

**Authors:** Mélanie Moreau, Anouk Chatefau, Jean-William Dupuy, Fabrice Cordelières, Violaine Moreau, Anne-Aurélie Raymond, Frédéric Saltel

## Abstract

Metastatic dissemination requires tumour cells to degrade the extracellular matrix through invadosomes, actin-rich degradative structures whose architecture adapts to the microenvironment. Here we identify the AAA+ ATPase RUVBL2 as a dual regulator of invadosome biology. RUVBL2 is enriched in tumours and accumulates dynamically within rosette invadosomes together with the R2TP components. RUVBL2 depletion abolishes invadosome formation and matrix degradation, reverts invasive cells to a stress-fibre phenotype and suppresses activating Src phosphorylation. Conversely, RUVBL2 overexpression induces degradation-competent invadosomes in non-invasive cells. This pro-invasive function does not require RUVBL2 ATPase activity: pharmacological or genetic ATPase inactivation preserves punctate invadosomes and matrix degradation but selectively abolishes rosette architecture, which instead depends on ATPase-driven assembly of an R2TP-like scaffold. RUVBL2 protein is elevated in tumours; its transcript levels are increased at metastatic stages in m and associated with poor survival in several cancers. RUVBL2 therefore couples the Src-dependent initiation of invasion to the higher-order structural organisation of invadosomes.

## Introduction

Metastasis remains the primary cause of cancer-related mortality. The dissemination of tumour cells relies on their capacity to degrade the extracellular matrix (ECM) and to migrate through adjacent tissues in order to invade and colonise distant organs. This degradative capacity is mediated by specialised F-actin-based structures termed invadosomes, which orchestrate the focal release of proteolytic enzymes including matrix metalloproteinases (MMPs) (Linder et al., 2023). Invadosomes are dynamic membrane protrusions that both engage ECM components and enzymatically degrade them (Paterson and Courtneidge, 2018). They display remarkable plasticity and adapt to their matrix environment: depending on cellular context and stimuli, they adopt distinct organisations ranging from individual dots and rosettes (circular assemblies) to linear structures aligned along collagen fibres (Di Martino et al., 2016; Ezzoukhry et al., 2016).

Invadosomes are sophisticated protein macrocomplexes that coordinate actin polymerisation through regulators such as Arp2/3, N-WASP, cortactin and dynamin, together with structural and scaffolding proteins including TKS5 (Seals et al., 2005). They also incorporate adhesion-associated proteins such as vinculin, talin, paxillin and matrix receptors like integrins, CD44 and discoidin domain receptors (DDRs) (Ezzoukhry et al., 2016; Linder et al., 2023). Their formation and dynamics are controlled by Rho-family GTPases, including CDC42, and by several kinases, most notably Src (Burger et al., 2014). Src occupies a central position in invadosome biology across cellular models: invadosomes were first identified in cells transformed by the v-Src oncogene (David-Pfeuty and Singer, 1980), and expression of constitutively active Src variants such as SrcY527F, used to generate the NIH3T3-Src cell line, remains a classical experimental model for invadosome studies (Seals et al., 2005). Invadosome-forming capacity correlates with metastatic potential (Leong et al., 2014), invadosomes have been documented in vivo within tumour microenvironments (Eckert et al., 2011; Gligorijevic et al., 2012; Lohmer et al., 2014) and they are required for cancer cell extravasation during metastatic dissemination (Leong et al., 2014). These structures therefore represent attractive therapeutic targets in oncology (Tsai, 2025).

To advance the molecular characterisation of invadosomes and to identify candidate targets within them, we previously profiled the invadosome proteome in the NIH3T3-Src model. Combining laser capture microdissection, which preserves molecular integrity, with mass spectrometry (LC-MS/MS), we identified 312 proteins enriched in invadosomes and revealed that rosette invadosomes possess their own active translation pole (Ezzoukhry et al., 2018). This last point was subsequently confirmed and characterised at the molecular level (Bonnard et al., 2026). Moreover, among the enriched proteins, RUVBL2 (Reptin) and its paralogue RUVBL1 (Pontin) displayed high enrichment ratios, suggesting a role in invadosome biology.

RUVBL2 is a highly conserved member of the AAA+ ATPase family involved in chromatin remodelling, transcriptional regulation, telomerase assembly and activity, DNA repair and senescence (Grigoletto et al., 2011; Raymond et al., 2015). In cancer, RUVBL2 has been implicated in tumour progression through multiple mechanisms (Tarangelo et al., 2015). It promotes cell proliferation in the intestinal epithelium and in colorectal cancer (Maurizy et al., 2021), drives hepatocellular carcinoma invasion by regulating Mep1a expression (Breig et al., 2017), contributes to epithelial-to-mesenchymal transition through mTOR pathway activation (Mao and Houry, 2017), supports lineage-specific transcriptional programmes in pancreatic cancer (Cunniff et al., 2025) and is essential for the oncogenic functions of MYC (Vogt et al., 2024). RUVBL2 is overexpressed in multiple tumour types, including head and neck squamous cell carcinoma (HNSCC), hepatocellular carcinoma (HCC) and lung adenocarcinoma (LUAD), where its expression correlates with poor prognosis (Su et al., 2022).

RUVBL2 also functions as a cytoplasmic co-chaperone within the R2TP complex, which is essential for the assembly and conformational maturation of numerous protein complexes (Lynham and Houry, 2022). The R2TP complex comprises four core subunits: RUVBL1, RUVBL2, RPAP3 and PIH1D1 (Lynham and Houry, 2022; Martino et al., 2018). It orchestrates the formation of macromolecular assemblies required for small nucleolar ribonucleoprotein (snoRNP) biogenesis, RNA polymerase II assembly and the stability of phosphatidylinositol 3-kinase-related protein kinases (PIKKs) (Lynham and Houry, 2022; Rivera-Calzada et al., 2017). Despite possessing conserved Walker A and Walker B motifs, RUVBL2 exhibits weak intrinsic catalytic activity as a monomer or homo-oligomer; integration into the R2TP complex markedly enhances its ATPase function, indicating that protein partners regulate its catalytic activity (Gorynia et al., 2011). Beyond the canonical R2TP, RUVBL2 participates in structurally related “R2TP-like” complexes that incorporate alternative adaptors, diversifying substrate recognition while preserving the fundamental mechanism of ATP hydrolysis (Maurizy et al., 2018; Rodríguez and Llorca, 2020; Santo et al., 2026).

Despite these well-characterised functions and the established role of RUVBL2 in cancer cell invasion, whether RUVBL2 is present and functional within invadosomes had not been addressed prior to our proteomic analysis. Based on these data, we hypothesised that RUVBL2 acts within an R2TP or R2TP-like complex to regulate the formation and activity of these structures. Here we show that RUVBL2 is both necessary and sufficient for invadosome formation through the control of Src activating phosphorylation, and that its ATPase activity is dispensable for invadosome formation and matrix degradation but strictly required for the higher-order rosette architecture.

## Results

### RUVBL2 is overexpressed in human tumours and localises to rosette invadosomes

To establish the clinical relevance of RUVBL2 in tumour invasion, we interrogated the CPTAC proteomic and TCGA transcriptomic databases across five cancer types: HNSCC, HCC, LUAD, colorectal adenocarcinoma (COAD) and renal cell carcinoma (RCC). RUVBL2 protein levels were significantly higher in tumour than in non-tumour samples in all five cancer types (Figure 1A). Comparing early-stage (stage I, localised) with advanced-stage (stage IV, metastatic) disease, RUVBL2 transcript levels were significantly increased at metastatic stages of HNSCC, HCC and RCC (Figure 1B). High RUVBL2 transcript levels were associated with reduced overall survival in HNSCC, HCC, LUAD and RCC (Figure S1). RUVBL2 expression therefore correlates with tumour progression and, specifically, with the metastatic transition.

**Figure 1.**
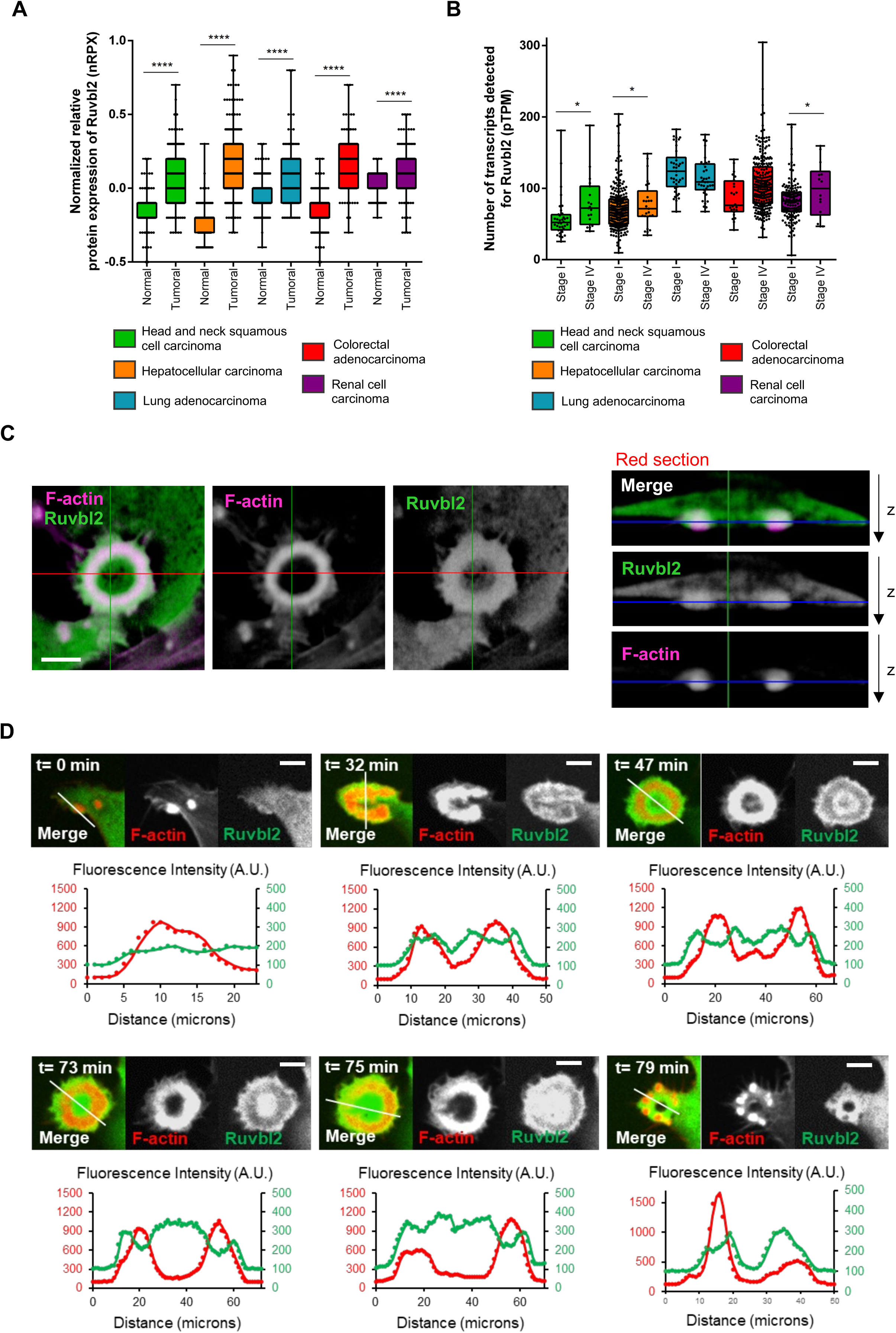
RUVBL2 is overexpressed in tumours, increased at metastatic stages, and localises to rosette invadosomes. (A) Relative RUVBL2 protein expression in head and neck squamous cell carcinoma (HNSCC), hepatocellular carcinoma (HCC), lung adenocarcinoma (LUAD), colorectal adenocarcinoma (COAD) and renal cell carcinoma (RCC) compared with matched non-tumour tissue, from the CPTAC database (ANOVA; p < 0.001; *p < 0.0001). (B) RUVBL2 transcript levels in stage I (localised) and stage IV (metastatic) disease for the same cancer types, from the TCGA database (ANOVA; p < 0.001; *p < 0.0001). (C) Localisation of RUVBL2-GFP in rosette invadosomes of living NIH3T3-Src cells expressing LifeAct-mCherry, imaged by lattice light-sheet microscopy. Scale bar, 5 µm. (D) Time-course analysis of RUVBL2-GFP localisation during rosette invadosome formation, maturation and disassembly by video-microscopy. Curves show fluorescence intensity within the selected region for LifeAct-mCherry (red) and RUVBL2-GFP (green). Scale bar, 5 µm.

We next examined whether RUVBL2 is present within invadosomes, as suggested by our previous proteomic analysis of the NIH3T3-Src model (Ezzoukhry et al., 2018). Three-dimensional lattice light-sheet imaging of NIH3T3-Src cells co-expressing LifeAct-mCherry and RUVBL2-GFP showed that RUVBL2 localises to rosette invadosomes, with additional enrichment in the intracellular region surrounding the actin-positive structures (Figure 1C).

Live-cell video-microscopy of the same model revealed a dynamic, stage-specific localisation during invadosome biogenesis (Figure 1D and Video S1). RUVBL2 accumulation became detectable only after the initial appearance of the actin dot (t = 0 min). During rosette formation, RUVBL2 localised within the invadosome with prominent enrichment at the periphery of the actin core (t = 32 and 47 min). Upon rosette maturation, RUVBL2 was enriched at both the periphery and the centre of the actin structure (t = 73 and 75 min). Finally, during rosette disassembly, RUVBL2 distribution became disorganised (t = 79 min). RUVBL2 was also detected in rosette invadosomes of two additional models, RAW 264.7 cells treated with RANK-L and HUVECs treated with VEGF (Figure S2A), indicating that its recruitment is not restricted to Src-transformed fibroblasts.

To determine whether the R2TP complex as a whole is recruited, we performed immunofluorescence analysis of fixed NIH3T3-Src cells expressing tagged versions of each subunit: RUVBL2-Flag, PIH1D1-DDK, RUVBL1-HA or RPAP3-DDK. All four components were detected in invadosomes, with particularly strong colocalisation between RUVBL2 and PIH1D1; both proteins labelled the entire invadosome but showed highest intensity at the outer edge (Figure S2B). Live-cell imaging of cells co-expressing LifeAct-mCherry and PIH1D1-GFP revealed dynamic localisation patterns closely similar to those of RUVBL2 (Figure S2C).

Collectively, these results establish that RUVBL2, together with at least one other R2TP component, is dynamically recruited to rosette invadosomes in several cellular models.

### RUVBL2 is required for invadosome formation and matrix degradation

To determine whether RUVBL2 is required for invadosome formation and activity, we used siRNA-mediated knockdown in the NIH3T3-Src model with two independent sequences.

RUVBL2 depletion markedly reduced both invadosome formation and matrix degradation (Figure 2A). This functional impairment was accompanied by an extensive reorganisation of the actin cytoskeleton: RUVBL2 knockdown not only decreased invadosome number but also significantly increased stress fibre formation, corresponding to a reversion towards the NIH3T3 wild-type (WT) phenotype (Figure 2B). The number of stress fibres per cell following RUVBL2 depletion approached the levels observed in NIH3T3-WT cells. Analysis of vinculin distribution showed that siRUVBL2 transfection promoted the formation of focal adhesions in place of the invadosomes typically observed in NIH3T3-Src cells, although relative vinculin intensity remained higher than in NIH3T3-WT cells (Figure 2B).

**Figure 2.**
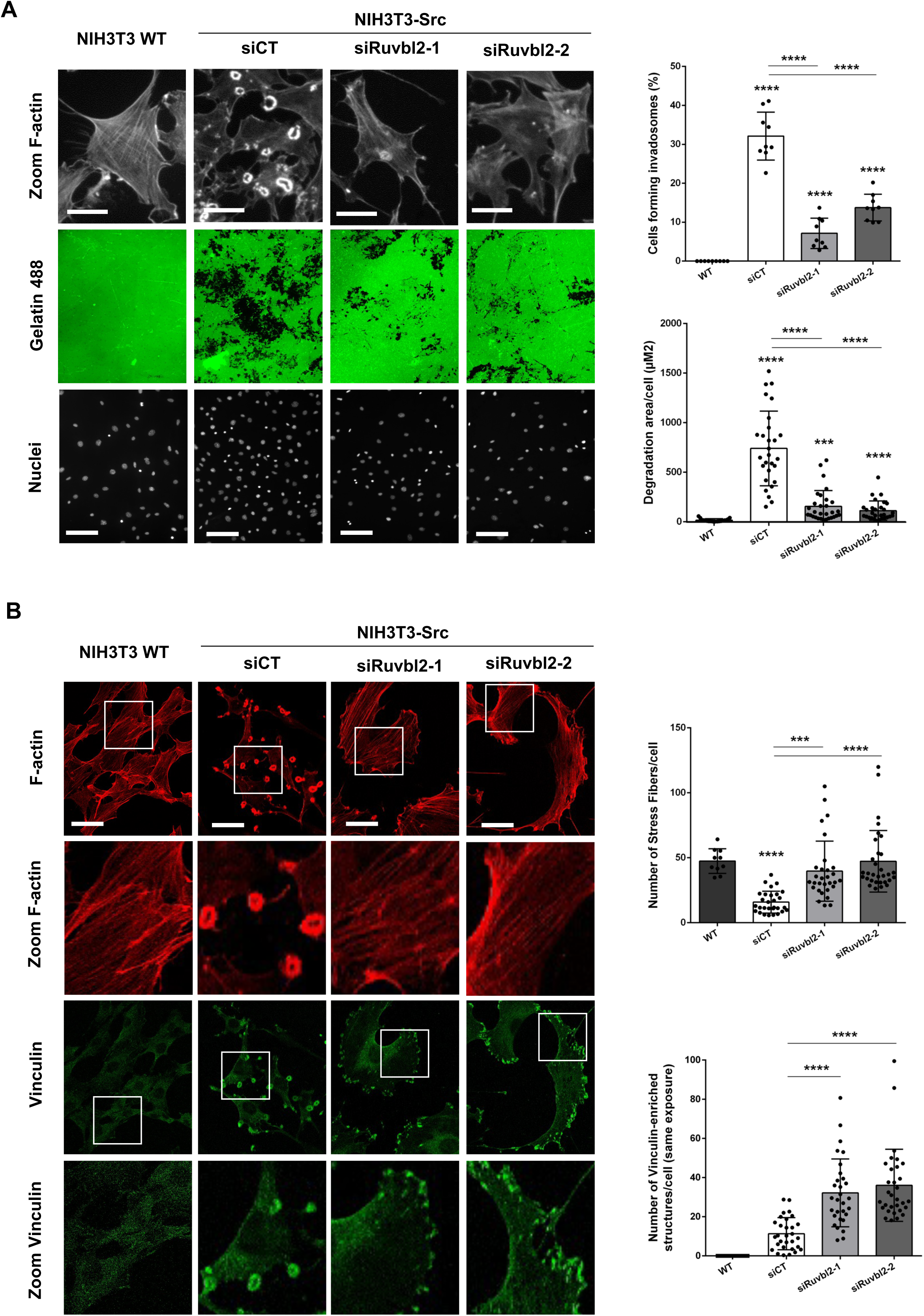
RUVBL2 is required for invadosome-mediated degradation and controls actin cytoskeleton organization. (A) Effect of RUVBL2 depletion on invadosome formation and degradative activity, measured by in situ zymography in NIH3T3-WT and NIH3T3-Src cells with or without RUVBL2 depletion. Quantification, right. Scale bar, 150 µm (n = 3; t test; ****p < 0.0001). (B) Effect of RUVBL2 depletion on actin cytoskeleton organisation, showing reversion to a WT-like phenotype with fewer rosette invadosomes and more stress fibres (SF). Upper panels, F-actin labelling alone and after processing with the FSegment algorithm for SF quantification (right). Lower panels, vinculin relocalisation from rosettes to focal adhesions in RUVBL2-depleted NIH3T3-Src cells. Scale bar, 30 µm. Quantification, right (n = 3; t test; p < 0.001; *p < 0.0001).

RUVBL2 depletion also reduced the levels of the other R2TP subunits (Figure S3A), consistent with the known interdependence of the complex, and PIH1D1 knockdown phenocopied RUVBL2 depletion with respect to invadosome formation and degradation (Figures S3B and S3C). These observations indicate that the R2TP module is coordinately regulated and that the phenotype we describe reflects the loss of a RUVBL2-containing complex rather than of RUVBL2 in isolation.

The reversion of NIH3T3-Src cells to a NIH3T3-WT-like cytoskeletal organisation upon RUVBL2 depletion suggested that RUVBL2 may modulate Src kinase activity or its downstream signalling.

### RUVBL2 sustains Src activating phosphorylation and the invadosome proteome

We first confirmed that Src and RUVBL2 colocalise within invadosomes (Figure 3A). We then measured Src expression and phosphorylation upon RUVBL2 depletion. Reduced RUVBL2 expression led to a marked decrease in activating phosphorylation of Src (pSrc Tyr416) without affecting total Src levels (Figure 3B).

**Figure 3.**
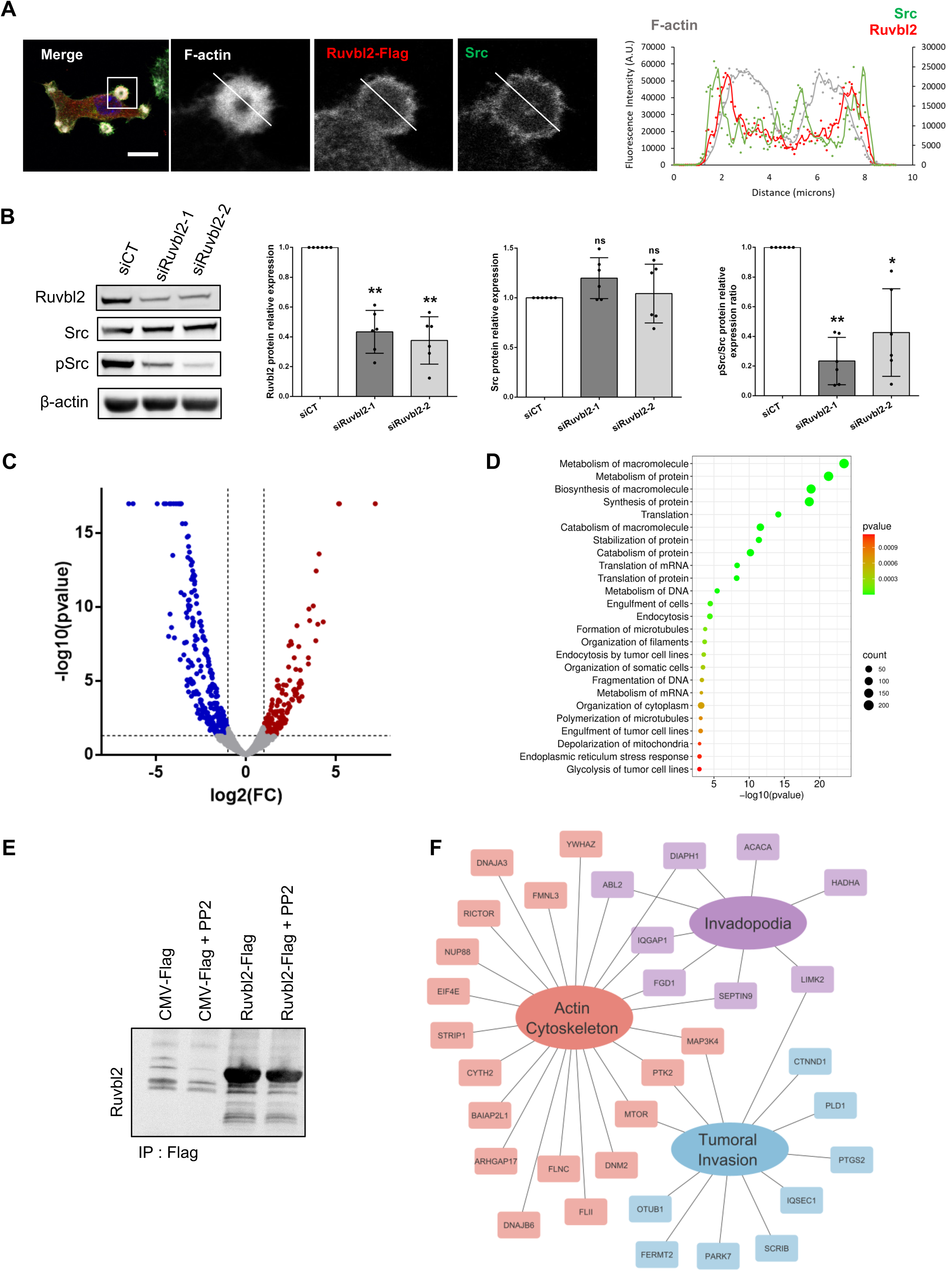
RUVBL2 depletion deregulates the Src pathway and the invadosome proteome. (A) Localisation of RUVBL2-Flag and endogenous Src at the invadosome periphery. Right, fluorescence intensity profile along the white line. Scale bar, 20 µm. (B) Effect of RUVBL2 depletion on Src phosphorylation. Western blot analysis of RUVBL2, Src and pSrc in NIH3T3-Src cells transfected with siRUVBL2. Quantification, right (n = 3; t test; *p < 0.05; **p < 0.01). (C) Mass spectrometry analysis of siRUVBL2-1-treated cells. The blue population represents proteins significantly decreased in siRUVBL2 relative to siCT. (D) Functional enrichment analysis of this protein set, incorporating expression ratios to determine Z-scores for the highlighted pathways. (E) Mass spectrometry analysis of co-immunoprecipitated RUVBL2-Flag samples from cells treated or not with 10 µM PP2 overnight. The red population represents proteins with a fold change (IP RUVBL2 versus IP RUVBL2 + PP2) ≥ 2 in all three replicates. (F) Supervised clustering of this protein set into the categories tumour invasion, invadosomes, actin cytoskeleton, R2TP members and protein turnover.

To evaluate the consequences of RUVBL2 depletion on invadosome components, we combined two approaches: quantification of established invadosome markers and Src regulators, and profiling of kinase activity using a PamGene PTK array. Protein levels of TKS5, cortactin, vinculin, dynamin 2 and PTP1B were unaffected, whereas MMP14, DDR2 and FAK1 were significantly decreased (Figure S4A). The mRNA levels of two Src regulators, Ptk2 (encoding FAK1) and Ptpn1 (encoding PTP1B), were unchanged (Figure S4B), indicating post-transcriptional control of FAK1 protein levels. Kinase activity profiling showed that the activity of 19 kinases belonging to the Src network was reduced upon RUVBL2 depletion, with Src itself being the most strongly affected, followed by FYN and PTK2B (FAK2); in contrast, MAPK1 activity was increased (Figures S4C and S4D). Src is therefore the kinase most sensitive to RUVBL2 loss within this network.

In parallel, we performed global proteomic analysis by mass spectrometry under RUVBL2 depletion, which identified 466 significantly deregulated proteins (Figure 3C and Table S1). Unsupervised functional enrichment revealed a major impact on protein metabolism and turnover, together with an effect on the cytoskeleton through the deregulation of microtubule formation, filament organisation and microtubule polymerisation (Figure 3D).

To further dissect the relationship between Src and RUVBL2, we characterised RUVBL2 binding partners under conditions of Src activation, which is constitutive in the NIH3T3-Src model. Immunoprecipitation of RUVBL2-Flag with or without PP2 treatment (an inhibitor of Src activating phosphorylation) identified 483 proteins whose interaction with RUVBL2 is reduced when Src activation is inhibited (Figures 3E and S5A and Table S2). Supervised clustering showed that a subset of these proteins is related to tumour invasion, actin cytoskeleton organisation and invadosomes (Figure 3F), functions consistent with the role of RUVBL2 within invadosomes, where it is enriched when Src is active. This analysis also revealed a substantial loss of interaction with known R2TP network members (Table 1) and with 292 proteins involved in protein metabolism. We confirmed the contribution of RUVBL2 to protein turnover by showing that RUVBL2 depletion significantly reduces global protein synthesis (Figure S5B).

**Table 1.** R2TP network members showing loss of interaction with RUVBL2 upon Src inhibition.

Together, these results indicate that RUVBL2 acts as an upstream regulator of invadosome formation through a dual input: it sustains Src activation and the activity of its downstream kinase network, and it supports the protein turnover and cytoskeletal organisation required for invadosome assembly.

### RUVBL2 is sufficient to induce invadosome formation in non-invasive cells

Given that RUVBL2 is required for invadosome formation and for Src activating phosphorylation, we asked whether RUVBL2 overexpression is sufficient to induce invadosomes in cells that do not normally form them.

We first expressed RUVBL2-Flag in NIH3T3-WT cells, which do not express the constitutively active form of Src. RUVBL2 overexpression significantly enhanced matrix degradation, and the actin structures formed were associated with gelatin degradation (Figure 4A). The structures responsible for this activity (Figure 4A, bottom panel) displayed a characteristic rosette morphology and showed enrichment of cortactin and phosphorylated Src (Figure 4B). At the level of the whole cell population, however, Src phosphorylation measured by western blot was not significantly changed (Figure 4C). Because only 5% of cells were transfected and 3% of transfected cells formed rosettes, this discrepancy most likely reflects dilution of the responding population in bulk lysates (whereas matrix degradation was quantified in transfected cells only) rather than absence of Src activation, as indicated by the clear pSrc enrichment detected at individual structures.

**Figure 4.**
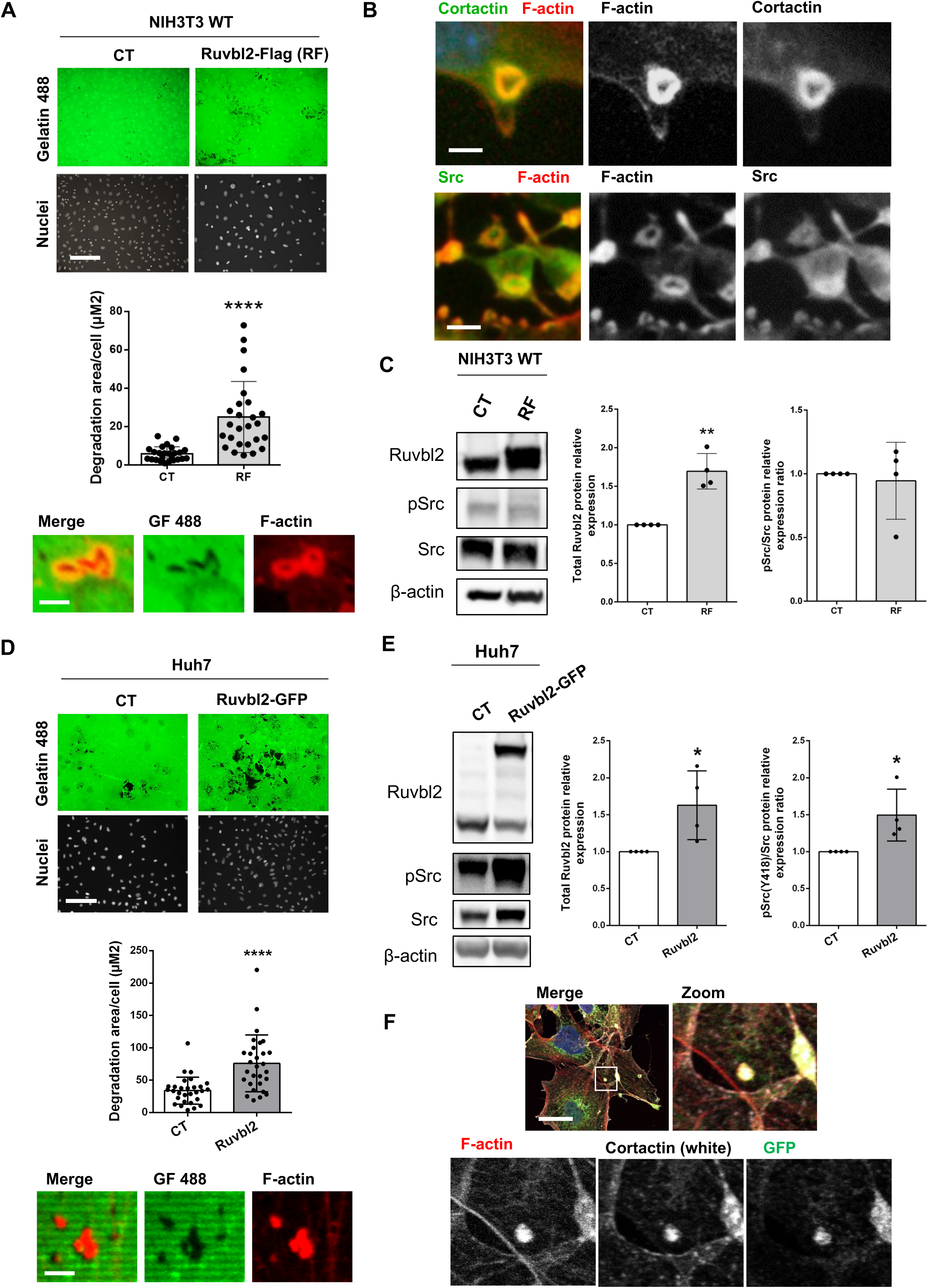
RUVBL2 overexpression is sufficient to induce invadosome formation in non-invasive cells. (A) In situ zymography of RUVBL2-overexpressing NIH3T3-WT cells. Scale bar, 150 µm. Quantification, middle; bottom, magnified views of degradative structures, scale bar 10 µm (n = 3; t test; ****p < 0.0001). (B) Colocalisation of cortactin and phosphorylated Src with F-actin in invadosomes induced by RUVBL2 expression. Scale bar, 10 µm. (C) Western blot analysis of RUVBL2, Src and pSrc in NIH3T3-WT cells transfected with CMV-Flag (CT) or RUVBL2-Flag (RF). Quantification, right (n = 3; t test; **p < 0.01). (D) In situ zymography of RUVBL2-overexpressing Huh7 cells. Scale bar, 150 µm. Quantification, middle (n = 3; t test; ****p < 0.0001); bottom, magnified views of degradative structures, scale bar 10 µm. (E) Western blot analysis of RUVBL2, Src and pSrc in Huh7 cells transduced with GFP alone (CT) or RUVBL2-GFP. Quantification, right (n = 3; t test; *p < 0.05). (F) Localisation of RUVBL2 at punctate invadosomes of Huh7 cells, colocalised with F-actin and cortactin. Scale bar, 30 µm.

We next used Huh7 cells, an HCC-derived line that is non-invasive under basal conditions (Breig et al., 2017). RUVBL2 overexpression in Huh7 cells induced invadosome formation and increased matrix degradation (Figure 4D) and concomitantly promoted Src activation, which was detectable at the population level in this model (Figure 4E). Notably, the invadosomes induced in Huh7 cells colocalised with cortactin and presented as large punctate structures rather than rosettes (Figure 4D, bottom panel, and Figure 4F).

Collectively, these findings demonstrate that RUVBL2 overexpression is sufficient to confer a degradative, invadosome-based phenotype on non-invasive cells, through Src relocalisation and activation. The punctate morphology of the induced structures in Huh7 further suggests that the acquisition of rosette architecture requires an additional, separable input.

### RUVBL2 ATPase activity dictates rosette architecture but is dispensable for invadosome formation

RUVBL2 lacks intrinsic kinase activity and therefore regulates Src indirectly. We next asked whether its ATPase activity contributes to invadosome formation, to Src activation, or to invadosome architecture. We used two complementary strategies: treatment with CB-6644, an allosteric inhibitor of RUVBL1-RUVBL2 ATPase activity (García-Martín et al., 2024), and expression of the ATPase-dead RUVBL2 mutant D299N (Grigoletto et al., 2013).

We first validated CB-6644 efficacy using mTOR levels as a readout of ATPase inhibition. CB-6644 treatment caused a partial but significant reduction in Src phosphorylation (Figure 5A) and a loss of rosette invadosomes (Figure 5B). Importantly, rosettes were replaced by large punctate invadosomes that retained matrix degradation capacity (Figure 5C). Thus, RUVBL1-RUVBL2 ATPase activity is dispensable for the formation and degradative function of punctate invadosomes but is required for rosette organisation.

**Figure 5.**
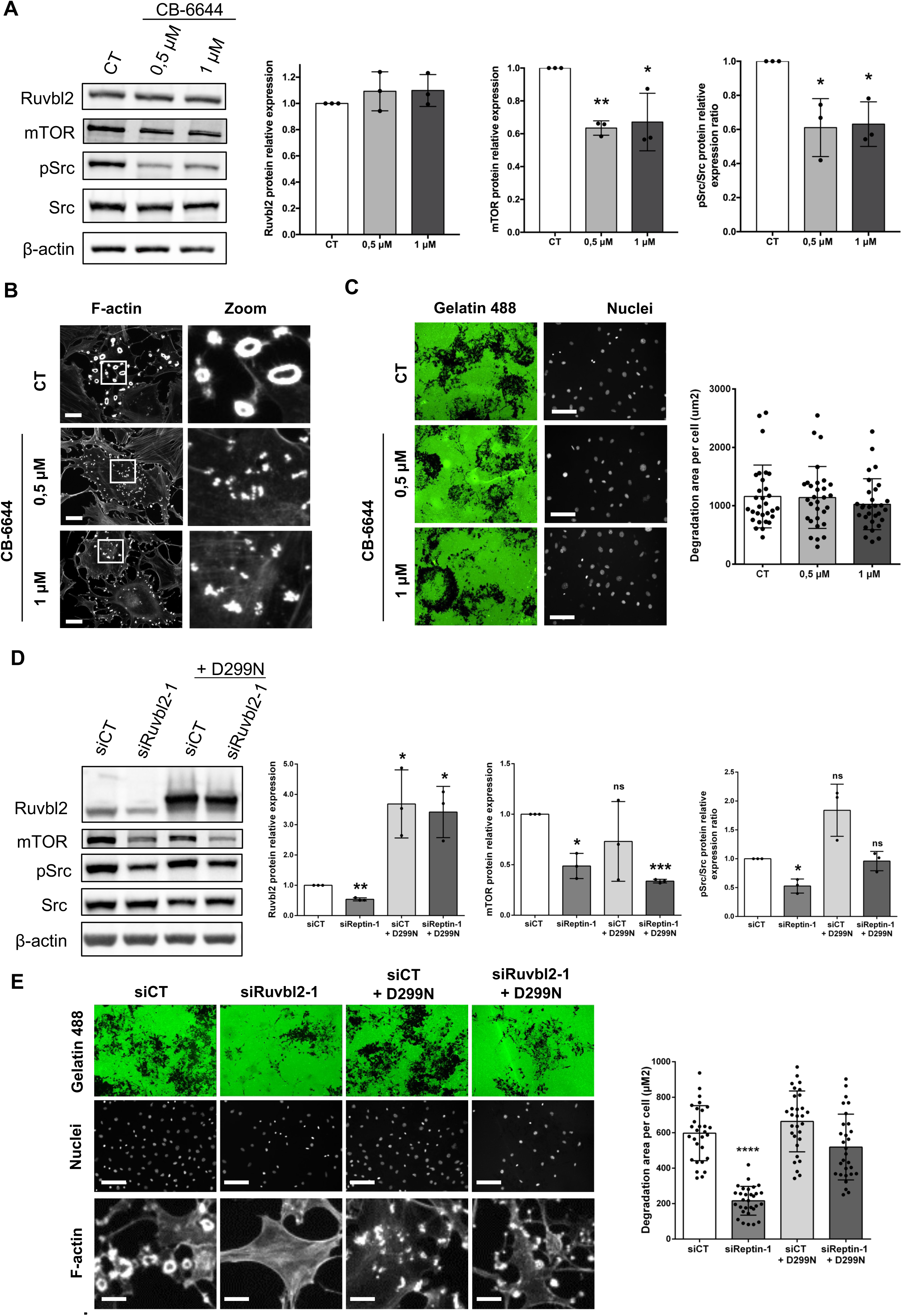
RUVBL2 ATPase activity determines rosette architecture but is dispensable for invadosome formation. (A) Western blot analysis of RUVBL2, mTOR, Src and pSrc in NIH3T3-Src cells treated with CB-6644 (0.5 and 1 µM). Quantification, right (n = 3; t test; *p < 0.05). (B) Views and magnification of actin structures in control and treated cells highlighting the rosette-to-punctate transition. Scale bar, 20 µm. (C) In situ zymography under the same conditions. Scale bar, 150 µm. Quantification, right. (D) Western blot analysis of RUVBL2, mTOR, Src and pSrc in NIH3T3-Src cells transfected with siRUVBL2-1 alone or co-transfected with the RUVBL2 D299N mutant. Quantification, right (n = 3; t test; p < 0.05; **p < 0.001). (E) In situ zymography under the same conditions. Scale bar, 150 µm. Insets, magnified views of actin structures, scale bar 20 µm. Quantification, right (n = 3; t test; ****p < 0.0001).

To attribute this effect specifically to RUVBL2 ATPase activity, we performed rescue experiments by expressing the siRUVBL2-1-resistant ATPase-dead D299N mutant in siRUVBL2-1-treated cells. This rescue restored both Src phosphorylation (Figure 5C) and matrix degradation (Figure 5D). However, the invadosomes formed under these conditions exhibited a large punctate morphology rather than the characteristic rosette architecture (Figure 5E), mimicking pharmacological ATPase inhibition.

The two approaches thus converge on invadosome architecture but differ with respect to Src: catalytically inactive RUVBL2 fully restores pSrc in depleted cells, whereas CB-6644 partially reduces it. This difference is expected, since CB-6644 inhibits both RUVBL1 and RUVBL2 and therefore affects the activity of the heterohexamer as a whole, whereas the D299N rescue restores RUVBL2 protein, and hence its scaffolding and co-chaperone functions, in a catalytically dead form. We therefore conclude that RUVBL2 protein, rather than its catalytic activity, is the principal determinant of Src activating phosphorylation, while ATPase activity provides an additional, quantitative level of control. Consistent with a partial contribution, residual degradative puncta persist under ATPase inhibition, whereas direct Src inhibition, for instance with PP2, is well established to abolish invadosome formation and matrix degradation.

Finally, we observed a decrease in PIH1D1 protein upon both pharmacological and genetic inhibition of RUVBL2 ATPase activity (Figure S6), consistent with destabilisation of the R2TP-like complex when its assembly is impaired.

Collectively, these results demonstrate that RUVBL2 ATPase activity is required for rosette invadosome architecture and contributes quantitatively to Src phosphorylation, but is not essential for the formation or degradative function of invadosomes.

## Discussion

Metastasis depends on the capacity of tumour cells to degrade the ECM through invadosomes, structures that coordinate proteolytic enzyme release and display considerable architectural plasticity. Our previous proteomic analysis of invadosomes in the NIH3T3-Src model (Ezzoukhry et al., 2018) identified RUVBL2 as highly enriched within these structures. Although RUVBL2 has been linked to tumour invasion and epithelial-to-mesenchymal transition, its role in invadosome biology had not been investigated. To our knowledge, this study is the first to demonstrate a direct mechanistic role for RUVBL2 in invadosome formation and function, and to separate two distinct activities of the protein: a catalysis-independent function that initiates invasion through Src, and an ATPase-dependent function that determines invadosome architecture.

Advanced imaging demonstrated that RUVBL2 is dynamically recruited to rosette invadosomes during their formation and maturation. Live-cell video-microscopy revealed a sequential pattern in which RUVBL2 becomes detectable after the initial actin dot appears, enriches at the periphery of the actin core during rosette formation and then distributes throughout peripheral and central regions upon maturation. All four R2TP components were detected within invadosomes, with particularly strong colocalisation between RUVBL2 and PIH1D1.

Functional characterisation showed that RUVBL2 depletion impairs both invadosome formation and matrix degradation, with extensive cytoskeletal reorganisation including increased stress fibres and the appearance of focal adhesions in place of invadosomes, effectively reverting NIH3T3-Src cells to a wild-type-like morphology. Mechanistically, RUVBL2 knockdown decreased Src activating phosphorylation without affecting total Src levels. A particularly informative aspect of this regulation arises from the nature of our model: NIH3T3-Src cells express a constitutively active Src bearing a mutation of the C-terminal inhibitory tyrosine (Tyr527). Because this regulatory residue can no longer be phosphorylated by CSK, Src is expected to remain in an open, autophosphorylation-competent conformation and to display constitutive activating phosphorylation at Tyr416. That RUVBL2 depletion markedly reduces this activating phosphorylation despite the inhibitory site being non-functional argues that RUVBL2 is required for the activating (auto)phosphorylation step itself, rather than acting through the canonical inhibitory branch of Src regulation.

As RUVBL2 lacks intrinsic kinase activity, it most likely promotes this activating phosphorylation indirectly. Several non-exclusive mechanisms can be envisaged: favouring the local clustering or scaffolding of Src molecules required for trans-autophosphorylation at Tyr416; sustaining the activity of Src-activating partners such as FAK; or assisting the conformational maturation of Src into a fully active kinase through its co-chaperone function. The last hypothesis is attractive given that Src is a well-characterised HSP90 client (Xu et al., 1999). The preservation of total Src levels argues against enhanced degradation and points instead to a defect in activation. We note, however, that our data establish RUVBL2 as required for Src activation but do not formally demonstrate that Src is the sole effector downstream of RUVBL2; epistasis experiments restoring Src activity in RUVBL2-depleted cells will be required to establish this point definitively.

Consistent with multi-level control of Src signalling, kinase activity profiling showed that RUVBL2 depletion reduced the activity of 19 kinases of the Src network, with Src itself most affected and including FAK2 (PTK2B). In line with this, FAK1 protein levels were significantly decreased upon RUVBL2 depletion, consistent with the established functional interdependence of FAK and Src at invadosomes: FAK autophosphorylation at Tyr397 creates a high-affinity binding site for Src, and this FAK–Src complex is critical for invadopodia dynamics and matrix degradation (Chan et al., 2009). Together with the preferential enrichment of RUVBL2 for actin- and invasion-related partners specifically when Src is active, these data support a self-reinforcing model in which Src activity promotes the assembly of RUVBL2-containing complexes, which in turn maintain Src activation.

Global proteomic analysis identified 466 deregulated proteins upon RUVBL2 depletion. Protein metabolism and turnover was the most strongly affected category, together with cytoskeletal organisation. This dual proteostasis/cytoskeleton signature was echoed by our interactome analysis: under Src activation, RUVBL2 preferentially associated with proteins linked to tumour invasion, actin cytoskeleton organisation and invadosomes, whereas Src inhibition triggered a massive loss of interaction with both canonical R2TP members and 292 proteins involved in protein metabolism.

The contribution of RUVBL2 to protein homeostasis is supported by converging observations. First, the decrease in FAK1 protein upon RUVBL2 depletion occurred without any change in the corresponding transcript, indicating post-transcriptional control of key invadosome effectors. Second, RUVBL2 depletion significantly reduced global protein synthesis. These results are consistent with the established function of the RUVBL1-RUVBL2/R2TP module as an HSP90 co-chaperone involved in the maturation and stabilisation of client proteins (Lynham and Houry, 2022; Muñoz-Hernández et al., 2019), and suggest that RUVBL2 sustains invadosome formation by maintaining the synthesis, maturation and stability of the proteins required for invadosome assembly and degradative activity.

Investigation of RUVBL2 enzymatic function using both pharmacological inhibition and an ATPase-dead mutant revealed that ATPase activity is specifically required for rosette architecture but dispensable for punctate invadosome formation and matrix degradation. This points to a model in which ATPase activity is required for R2TP-like complex assembly and structural stabilisation. ATPase activity is indeed essential for proper R2TP complex formation, client stabilisation and the structural integrity of the RUVBL1-RUVBL2 heterohexameric ring (Martino et al., 2018; Muñoz-Hernández et al., 2019). Cryo-EM studies have shown that PIH1D1 binding to the DII domain of RUVBL2 induces conformational rearrangements that regulate nucleotide exchange, coupling protein–protein interactions to ATPase activity (Muñoz-Hernández et al., 2019). Our observation of reduced PIH1D1 protein upon pharmacological and genetic ATPase inhibition supports this model. PIH1D1 stability is critically dependent on its incorporation into the R2TP complex, and PIH1D1 undergoes rapid degradation when not associated with its partners (Henri et al., 2018; Zhao et al., 2008); the diminished PIH1D1 levels we observe therefore most likely reflect destabilisation consequent to impaired complex assembly rather than transcriptional downregulation.

An apparent discrepancy deserves comment. CB-6644 reduced Src phosphorylation, whereas the ATPase-dead D299N mutant restored it in depleted cells. We interpret this as reflecting the distinct nature of the two perturbations: CB-6644 inhibits the catalytic activity of both RUVBL1 and RUVBL2 within the heterohexamer and may additionally affect RUVBL1-dependent functions, whereas the D299N rescue restores RUVBL2 protein, and therefore its scaffolding and co-chaperone roles, in a catalytically inactive form. The simplest model consistent with both observations is that the presence of RUVBL2 protein, rather than its catalytic cycle, is the principal requirement for Src activation, with ATPase activity providing an additional quantitative contribution. We cannot formally exclude a RUVBL1-specific contribution to Src regulation, and distinguishing the individual contributions of the two paralogues will require selective genetic approaches.

Gain-of-function experiments demonstrated that RUVBL2 overexpression induces invadosome formation in both NIH3T3-WT cells and the non-invasive HCC line Huh7. In NIH3T3-WT cells, RUVBL2 overexpression enhanced matrix degradation and promoted pSrc enrichment within newly formed rosettes, whereas in Huh7 cells it triggered both Src activation and the formation of degradation-competent punctate invadosomes. By demonstrating that RUVBL2 is sufficient to trigger invadosome formation in non-invasive cells, these experiments identify RUVBL2 as a potent inducer of the invasive phenotype.

Based on these findings, we propose a hierarchical model for RUVBL2 function in invadosome biology. In the absence of RUVBL2, cells cannot form invadosomes and revert to a non-invasive phenotype characterised by stress fibres and focal adhesions. RUVBL2 overexpression alone is sufficient to induce invadosome formation, even in non-invasive cells, by enabling the activating autophosphorylation of Src; the morphology of the structures induced under these conditions depends on the cellular context, and possibly on the basal activation state of Src: rosettes in NIH3T3-WT fibroblasts, large punctate invadosomes in Huh7 cells. The transition from punctate to rosette architecture requires the integration of RUVBL2 into a functional R2TP-like complex containing at least PIH1D1, as indicated by their strong colocalisation, similar recruitment dynamics and the loss of rosettes upon ATPase inhibition. We therefore propose that RUVBL2-mediated Src activation is sufficient to initiate invadosome formation, whereas the R2TP-like complex acts as a molecular scaffold essential for the higher-order organisation characteristic of mature rosettes.

This proposed scaffolding function is consistent with the established role of RUVBL1-RUVBL2 as a versatile platform for protein–protein interactions (Dauden et al., 2021). Cryo-EM studies of chromatin remodelling complexes such as INO80 have shown that the RUVBL1-RUVBL2 heterohexamer provides a structural scaffold organising the architecture of other subunits (Aramayo et al., 2018). Similarly, RUVBL1-RUVBL2 is essential for the assembly of the γ-tubulin ring complex (γTuRC), where it promotes the ordered assembly of multiple subunits without being incorporated into the final complex (Zimmermann et al., 2020). By analogy, we propose that an R2TP-like complex functions as an assembly platform coordinating the recruitment and spatial organisation of invadosome components required for rosette formation.

Recent work has identified alternative R2TP-like complexes containing distinct adaptors, such as R2SP (RUVBL1-RUVBL2-SPAG1-PIH1D2), which displays tissue-specific expression and substrate specificity (Maurizy et al., 2018; Santo et al., 2026). Whether similar specialised complexes operate in invadosome biology, potentially incorporating tissue-specific or context-dependent adaptors, warrants further investigation and may provide an additional regulatory layer across cell types and microenvironmental conditions.

Our analysis of CPTAC and TCGA datasets indicates that RUVBL2 protein levels are significantly elevated across five major cancer types, that elevated transcript levels correlate with reduced overall survival in four of them, and that transcript levels are increased at metastatic stages compared with localised disease in three of them (HNSCC, HCC and RCC). These clinical correlations, together with the demonstration that RUVBL2 is both necessary and sufficient for invadosome-mediated degradation, position RUVBL2 as a candidate therapeutic target for anti-metastatic strategies. The separability of its two functions is of particular interest in this context: ATPase inhibitors such as CB-6644 would be expected to suppress rosette architecture while leaving punctate, degradation-competent invadosomes intact, suggesting that targeting RUVBL2 protein levels or its scaffolding interfaces, rather than its catalytic activity alone, may be required to fully suppress invasion.

## Key resources table

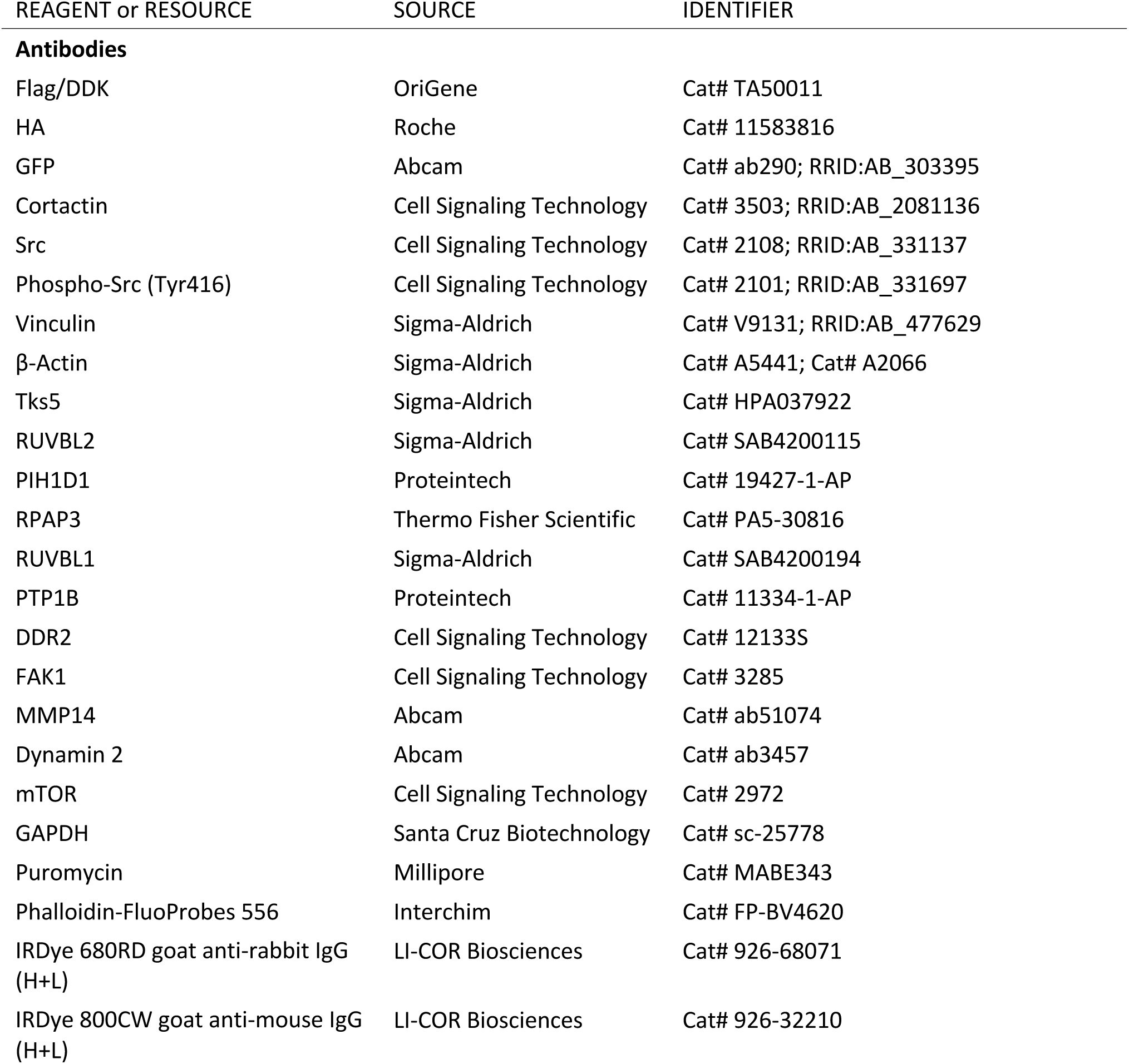
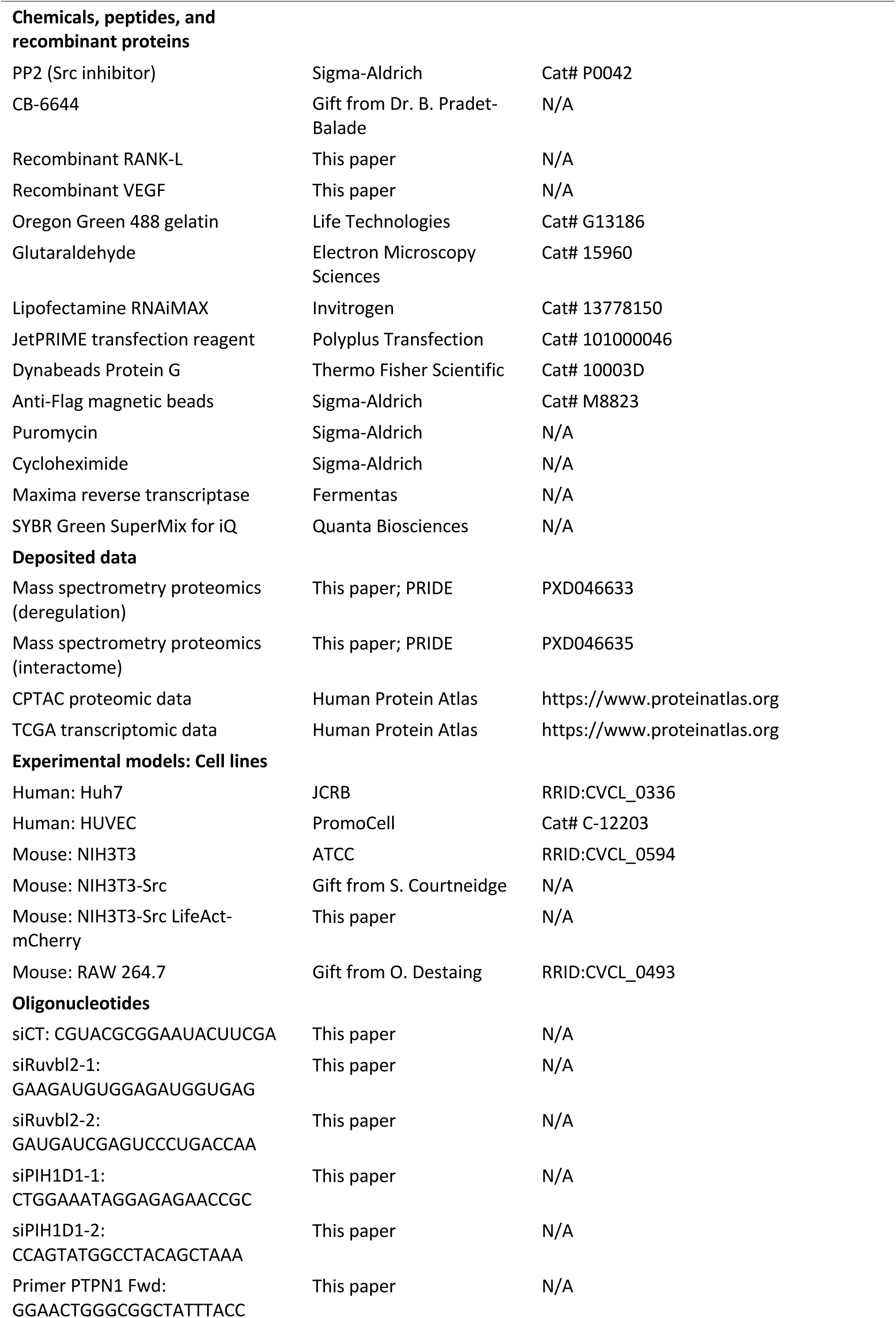
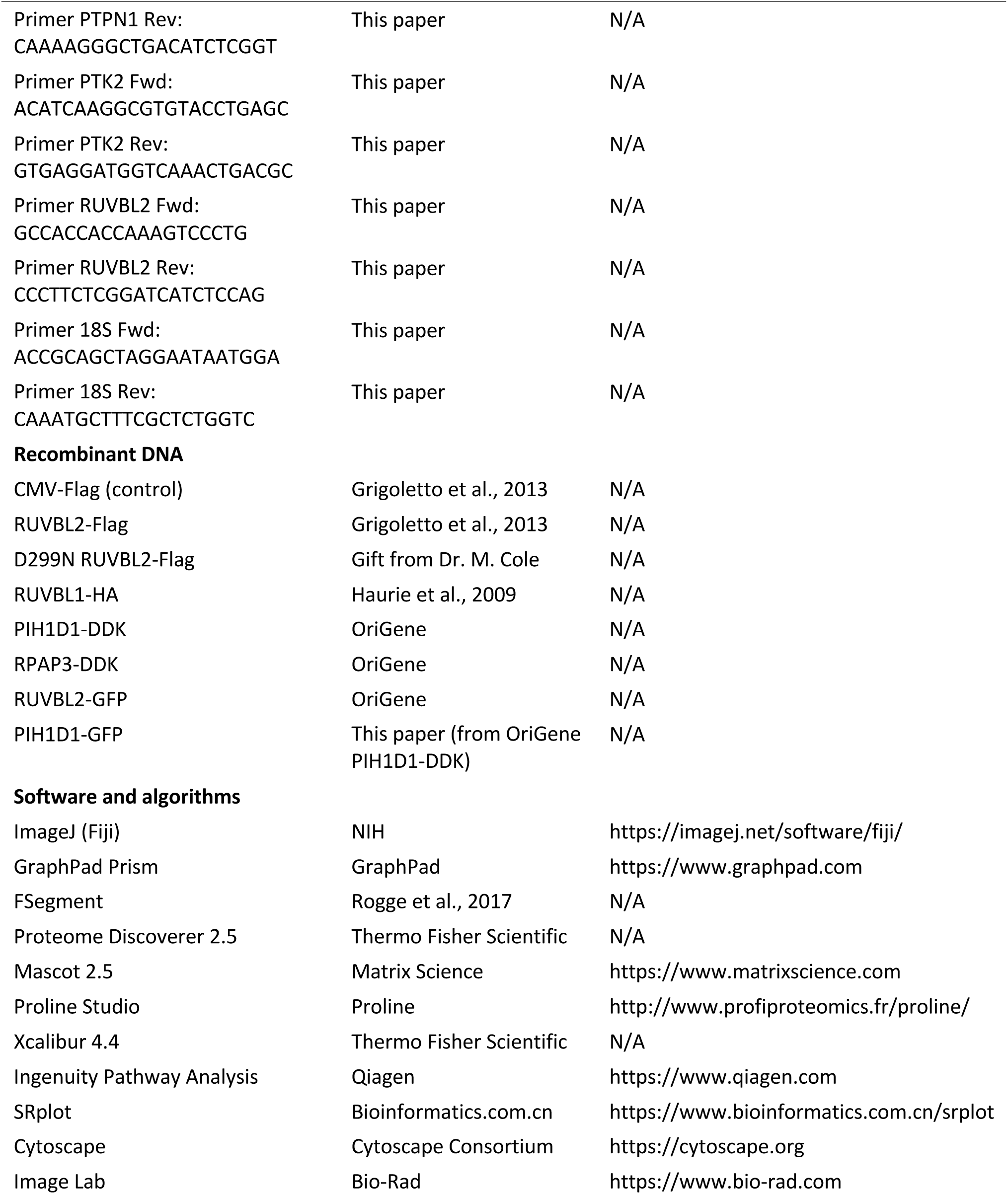

## Resource availability

### Lead contact

Further information and requests for resources and reagents should be directed to and will be fulfilled by the lead contact, Frédéric Saltel.

### Materials availability

Plasmids and cell lines generated in this study are available from the lead contact upon reasonable request. The CB-6644 compound and the D299N RUVBL2-Flag construct were obtained from third parties and are subject to their respective material transfer agreements.

### Data and code availability

- The mass spectrometry proteomics data have been deposited in the ProteomeXchange Consortium via the PRIDE partner repository (Deutsch et al., 2023; Perez-Riverol et al., 2025) with the dataset identifiers PXD046633 and PXD046635, and are publicly available as of the date of publication. Publicly available datasets analysed in this study were obtained from CPTAC and TCGA through the Human Protein Atlas. Original western blot images (source data) are provided with this paper.
- This paper does not report original code.
- Any additional information required to reanalyse the data reported in this paper is available from the lead contact upon request.

### Experimental model and study participant details

#### Cell lines

Huh7, HUVEC and NIH3T3 cell lines were obtained from JCRB, PromoCell and the American Type Culture Collection, respectively. NIH3T3-Src cells were a gift from S. Courtneidge (Burnham Institute for Medical Research); NIH3T3-Src LifeAct-mCherry cells are derived from this cell line; and RAW 264.7 cells were a donation from O. Destaing (Institute for Advanced Biosciences). Huh7, NIH3T3-Src, NIH3T3 and RAW 264.7 cells were cultured in Dulbecco’s modified Eagle’s medium with 4.5 g l−1 glucose and GlutaMAX-I (Gibco) supplemented with 10% fetal calf serum (EuroBio). HUVECs were cultured in Endothelial Cell Growth Medium 2 with SupplementMix (PromoCell, C22011). Cells were maintained at 37°C in a humidified atmosphere with 5% CO2 and confirmed to be free of mycoplasma by PCR.

For invadosome induction, RAW 264.7 cells were cultured in αMEM with 10% fetal calf serum (EuroBio) and treated with RANK-L at 50 ng ml−1 for 4 days, and HUVECs were treated with VEGF at 50 ng ml−1 for 24 h.

For inhibition experiments, adherent cells were treated with PP2 (P0042, Sigma) at 10 µM overnight, or with CB-6644 (kind gift from Dr. B. Pradet-Balade) at 0.5 and 1 µM for 24 h.

#### Method details

##### siRNA and plasmid transfection

Small interfering RNAs (siRNAs) were transfected using Lipofectamine RNAiMAX (Invitrogen) according to the manufacturer’s instructions. An siRNA targeting Gl2 luciferase was used as a negative control. Silencing efficiency was determined by western blotting. The siRNAs used were: siCT (CGUACGCGGAAUACUUCGA); siRuvbl2-1 (GAAGAUGUGGAGAUGGUGAG); siRuvbl2-2 (GAUGAUCGAGUCCCUGACCAA); siPIH1D1-1 (CTGGAAATAGGAGAGAACCGC) and siPIH1D1-2 (CCAGTATGGCCTACAGCTAAA).

Plasmids were transfected using JetPRIME (Polyplus Transfection) according to the manufacturer’s instructions. Transfection efficiency was assessed by western blot or immunofluorescence. Plasmids used were: CMV-Flag control (Grigoletto et al., 2013), RUVBL2-Flag (Grigoletto et al., 2013), D299N RUVBL2-Flag (gift from Dr. M. Cole), RUVBL1-HA (Haurie et al., 2009), PIH1D1-DDK (OriGene) and RPAP3-DDK (OriGene).

#### Lentiviral transduction

Cells were seeded in six-well plates at 105 cells per well. RUVBL2-GFP lentivirus (constructed from the OriGene RUVBL2-GFP plasmid), PIH1D1-GFP lentivirus (constructed from the OriGene PIH1D1-DDK plasmid) or control GFP lentivirus was added 7 h after cell seeding, at a multiplicity of infection of 5. The culture medium was replaced daily for three days, and cells were used after the final change.

##### In situ degradation assay

Glass coverslips were coated with fluorescent gelatin (Oregon Green gelatin, Life Technologies G13186) for 20 min at room temperature, fixed with 0.5% glutaraldehyde (Electron Microscopy Sciences 15960) for 40 min at room temperature and washed three times in PBS (Gibco). Cells were then seeded at 15,000 cells per well in 24-well plates and cultured for 24 h. Areas of degradation, which no longer fluoresce, were quantified using ImageJ.

### Immunofluorescence, video-microscopy and 3D microscopy

Cells were fixed with 4% paraformaldehyde (pH 7.2) for 10 min, permeabilised with 0.2% Triton X-100 for 10 min and incubated with the indicated antibodies. Imaging was performed on an Axioplan 2 (Zeiss) epifluorescence microscope for degradation assays, or on an SP5 confocal microscope (Leica) with a Plan-Neofluar objective for colocalisation. Each channel was imaged sequentially using the multitrack module before merging, to prevent bleed-through between fluorochromes. The following antibodies were used: Flag/DDK (OriGene, TA50011), HA (Roche, 11583816), GFP (Abcam, ab290), cortactin (Cell Signaling, 3503), Src (Cell Signaling, 2108) and vinculin (Sigma, V9131). F-actin was labelled with Phalloidin-FluoProbes 556 (Interchim, FP-BV4620). Live-cell video-microscopy was performed on a DMi8 microscope (Leica) at the Bordeaux Imaging Center, and three-dimensional imaging on a Lattice Lightsheet 7 (Zeiss).

### Stress fibre quantification

Stress fibre number and length were automatically quantified using the FSegment algorithm running on MATLAB, as described (Rogge et al., 2017).

### Immunoprecipitation

Cells were lysed in PBS with 1% Triton X-100 supplemented with protease and phosphatase inhibitors. One milligram of solubilised protein in 500 µl was incubated with primary antibody overnight at 4°C, followed by addition of 20 µl Dynabeads Protein G (Thermo Fisher, 10003D) for 1 h at 4°C; alternatively, lysates were incubated for 1 h with anti-Flag magnetic beads (Sigma). After three washes in lysis buffer, pellets were resuspended in Laemmli buffer for western blot and mass spectrometry analysis.

### Western blotting

Proteins were extracted from cell lysates in radio-immunoprecipitation assay buffer (25 mM Tris-HCl pH 7.5, 150 mM NaCl, 1% IGEPAL, 1% sodium deoxycholate, 0.1% SDS) supplemented with protease and phosphatase inhibitors. Cell lysates were centrifuged at 10,000 g for 15 min. Supernatants were denatured with Laemmli loading buffer containing 2.5% 2-β-mercaptoethanol at 95°C for 10 min and separated by SDS-PAGE on 10% acrylamide gels (Fast Cast kit, Bio-Rad). Proteins were transferred to nitrocellulose membranes (Trans-Blot Turbo midi-size, Bio-Rad), blocked with Odyssey blocking buffer (LI-COR) and probed with primary antibodies, diluted 1/1,000, overnight at 4°C. The following antibodies were used: β-actin (Sigma, A5441 and A2066), cortactin (Cell Signaling, 3503), vinculin (Sigma, V9131), Tks5 (Sigma, HPA037922), RUVBL2 (Sigma, SAB4200115), PIH1D1 (Proteintech, 19427-1-AP), RPAP3 (Thermo Fisher, PA5-30816), RUVBL1 (Sigma, SAB4200194), PTP1B (Proteintech, 11334-1-AP), DDR2 (Cell Signaling, 12133S), Src (Cell Signaling, 2108), pSrc (Cell Signaling, 2101), FAK1 (Cell Signaling, 3285), MMP14 (Abcam, ab51074), dynamin 2 (Abcam, ab3457), mTOR (Cell Signaling, 2972), GAPDH (Santa Cruz, sc-25778) and puromycin (Millipore, MABE343).

Primary antibodies were diluted in Tris-buffered saline (TBS) with 5% bovine serum albumin. After washing twice for 10 min in TBS with 0.1% Tween-20, membranes were incubated for 1 h with species-specific far-red fluorescent secondary antibodies: IRDye 680RD goat anti-rabbit IgG (H+L) (LI-COR) or IRDye 800CW goat anti-mouse IgG (H+L) (LI-COR), diluted 1:5,000 in TBS with 5% BSA. After two 10-min washes in TBS with 0.1% Tween-20 and a final wash in TBS, membranes were imaged on a ChemiDoc imager (Bio-Rad). Bands of the expected size were quantified with Image Lab software. Uncropped blots are provided in the source data file.

### RT-qPCR

Total RNA was extracted using the NucleoSpin RNA kit (Macherey-Nagel) according to the manufacturer’s instructions. cDNA was synthesised from 1 µg of total RNA with Maxima reverse transcriptase (Fermentas). Twenty nanograms of cDNA were amplified on a CFX96 Real-Time PCR detection system (Bio-Rad) using SYBR Green SuperMix for iQ (Quanta Biosciences) with the following cycling conditions: initial denaturation at 95°C for 10 min, followed by 40 cycles of 95°C for 15 s and 60°C for 1 min. Expression was normalised to 18S ribosomal RNA, and relative levels were calculated using the comparative (2−ΔΔCt) method. The following primers were used: PTPN1 (Fwd GGAACTGGGCGGCTATTTACC; Rev CAAAAGGGCTGACATCTCGGT), PTK2 (Fwd ACATCAAGGCGTGTACCTGAGC; Rev GTGAGGATGGTCAAACTGACGC), RUVBL2 (Fwd GCCACCACCAAAGTCCCTG; Rev CCCTTCTCGGATCATCTCCAG), 18S (Fwd ACCGCAGCTAGGAATAATGGA; Rev CAAATGCTTTCGCTCTGGTC).

### SUnSET assay

Cells were treated with 10 µg ml−1 puromycin for 10 min at 37°C before protein extraction and detection by western blotting. As a negative control, cells were pre-treated with 35 µM cycloheximide for 10 min at 37°C.

### Kinase activity profiling

Protein tyrosine kinase activity was profiled on PamChip arrays (PamGene), following the kit’s instructions, at the BioProt platform (Bordeaux Neurocampus). The acquisition and analysis were conducted by PamGene using company-specific analysis software.

### Mass spectrometry analysis

Sample preparation and trypsin digestion were performed as described (Henriet et al., 2017). NanoLC-MS/MS analysis was carried out on an Ultimate 3000 RSLC Nano-UHPLC system (Thermo Scientific) coupled to a nanospray Orbitrap Fusion Lumos Tribrid mass spectrometer (Thermo Fisher Scientific). Peptide extracts were loaded onto a 300 µm ID × 5 mm PepMap C18 precolumn (Thermo Scientific) at 10 µl min−1. After a 3-min desalting step, peptides were separated on a 50-cm EasySpray column (75 µm ID, 2 µm C18 beads, 100 Å pore size, ES903, Thermo Fisher Scientific) with a 4–40% linear gradient of solvent B (0.1% formic acid in 80% acetonitrile) over 57 min at 300 nl min−1. The mass spectrometer operated in positive ion mode at 2.0 kV needle voltage. Data were acquired with Xcalibur 4.4 in data-dependent mode. MS scans (m/z 375–1,500) were recorded at a resolution of 120,000 (at m/z 200) with a standard AGC target and automatic injection time, followed by a top-speed duty cycle of up to 3 s for MS/MS acquisition. Precursor ions (charge states 2–7) were isolated in the quadrupole with a 1.6 Th window and fragmented by HCD at 28% normalised collision energy. MS/MS spectra were acquired in the Orbitrap at a resolution of 30,000 (at m/z 200) with a standard AGC target and automatic maximum injection time. Selected precursors were excluded for 60 s.

Protein identification was performed in Proteome Discoverer 2.5 using the Mascot 2.5 algorithm against a UniProt Mus musculus database (55,338 entries, release January 2022). Two missed trypsin cleavages were allowed. Mass tolerances were 10 ppm in MS and 0.02 Da in MS/MS. Oxidation (M) and acetylation (K) were set as dynamic modifications and carbamidomethylation (C) as a static modification. Raw data were imported into Proline Studio for feature detection, alignment and quantification. Protein identification was accepted with at least two specific peptides at rank 1 and a protein FDR below 1%, calculated using the decoy option in Mascot. Label-free MS1 quantification by extracted ion chromatograms was performed with parameters described previously (Henriet et al., 2017). Normalisation was carried out on the median of ratios, and missing values were imputed at 5% of background noise.

For interactome analysis, RUVBL2-Flag immunoprecipitates from NIH3T3-Src cells treated or not with 10 µM PP2 overnight were analysed in three independent replicates. Src-dependent RUVBL2 partners were defined as proteins with a fold change (IP RUVBL2 versus IP RUVBL2 + PP2) ≥ 2 in all three replicates.

### Bioinformatics

Proteomic data were analysed with Ingenuity Pathway Analysis (Qiagen). Bioinformatic representations were generated with SRplot (https://www.bioinformatics.com.cn/srplot) and Cytoscape. CPTAC protein and TCGA transcriptomic data were accessed through the Human Protein Atlas (Thul and Lindskog, 2018).

### Quantification and statistical analysis

Data are presented as mean ± s.e.m. of at least three independent experiments. For comparisons between two conditions, two-tailed unpaired Student’s t-tests were used. For comparisons involving more than two conditions or more than one cancer type, one-way ANOVA followed by correction for multiple comparisons was applied. Analyses were performed in GraphPad Prism. Significance levels are indicated as p < 0.05; p < 0.01; p < 0.001; ****p < 0.0001. Exact p values, n numbers and the statistical test used are given in each figure legend. No statistical method was used to predetermine sample size, no data were excluded, experiments were not randomised and investigators were not blinded to allocation during experiments and outcome assessment.

## Acknowledgments

We thank La Ligue contre le Cancer (2023 Certified Team) and SIRIC BRIO for funding. We thank the Vect’UB platform (TBM Core US005) for technical support with lentiviral production, the OneCell platform (TBM Core US005) for support with qPCR analysis, the BioProt platform (Bordeaux Neurocampus) for the production and acquisition of PamChip data, and the Bordeaux Imaging Center for microscopy. We thank Elodie Darbo for help with Cytoscape, Dr. B. Pradet-Balade for the generous gift of CB-6644 and Dr. M. Cole for the D299N RUVBL2-Flag construct.

## Author contributions

M.M. performed the majority of experiments, participated in writing the manuscript, analysed data and prepared figures. A.C. contributed to cell biology experiments. J.-W.D. performed mass spectrometry acquisition and analysis. F.C. developed image analysis pipelines. V.M. contributed to experimental design and interpretation. A.-A.R. supervised the study. F.S. conceived and supervised the study, secured funding and wrote the manuscript with input from all authors.

## Supplemental information

Document S1. Figures S1–S6.

**Figure S1.**
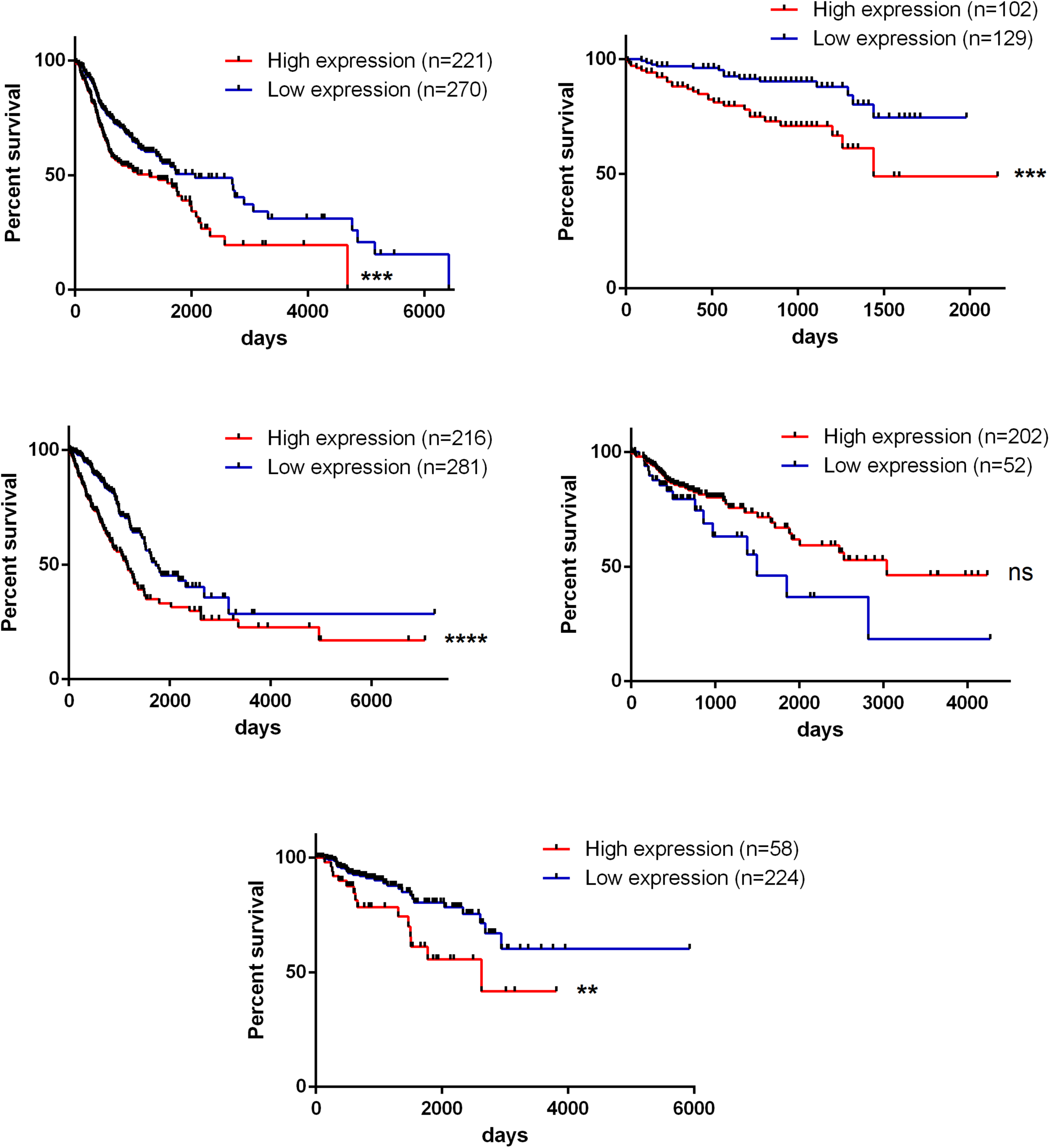
**High RUVBL2 mRNA expression correlates with poor prognosis**, related to Figure 1. Overall survival curves stratified by RUVBL2 mRNA expression in head and neck squamous cell carcinoma (A), hepatocellular carcinoma (B), lung adenocarcinoma (C), colorectal adenocarcinoma (D) and renal cell carcinoma (E) (log-rank test; ****p < 0.0001).

**Figure S2.**
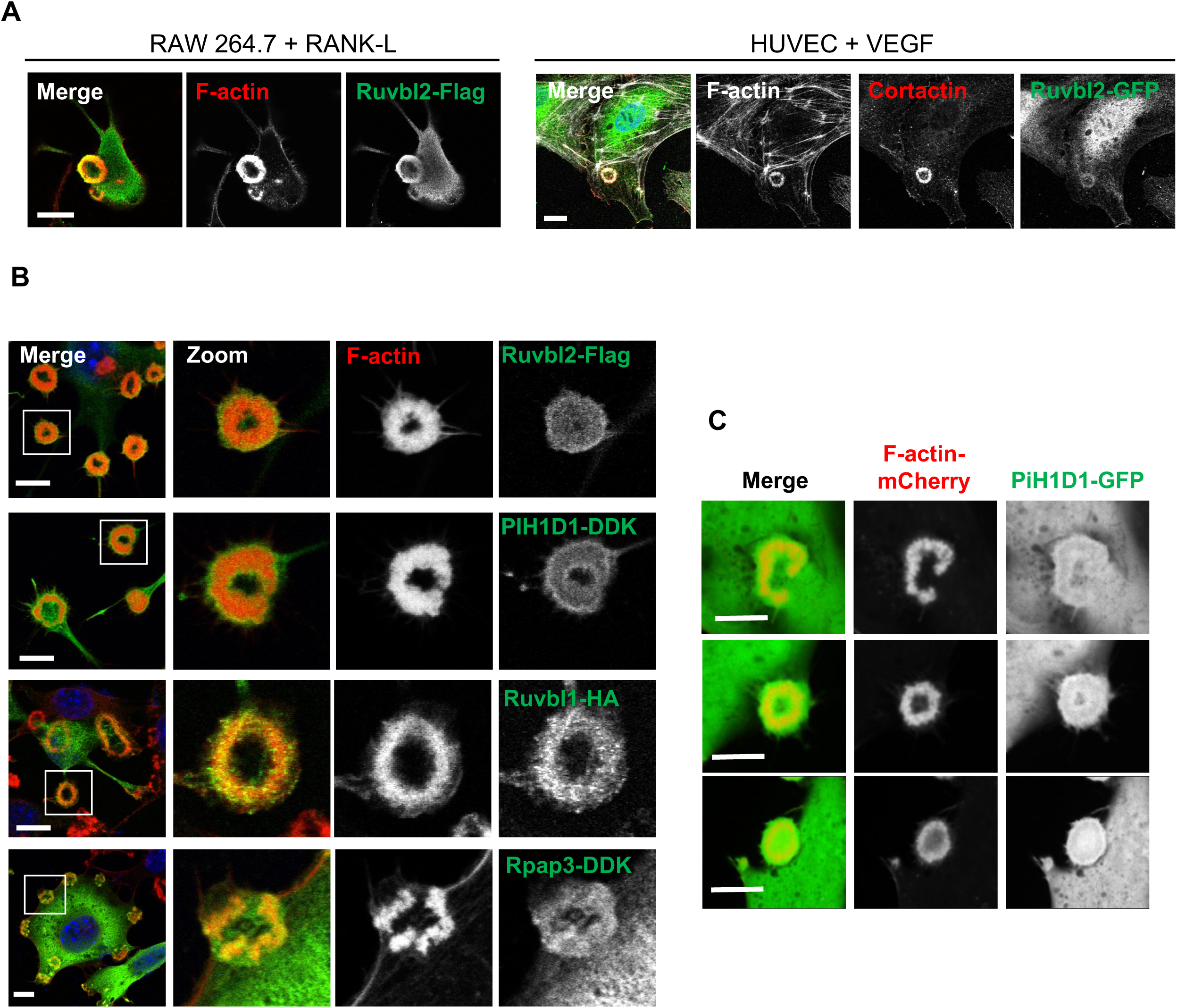
**RUVBL2 is present in different models of rosette invadosomes, and the R2TP complex localises to NIH3T3-Src invadosomes**, related to Figure 1. (A) Localisation of RUVBL2 in rosette invadosomes of RAW 264.7 cells treated with RANK-L and HUVECs treated with VEGF. Scale bar, 10 µm (n = 3; t test; p < 0.05; p < 0.01). (B) Localisation of tagged RUVBL2, PIH1D1, RUVBL1 and RPAP3 at NIH3T3-Src invadosomes (F-actin labelling). Scale bar, 10 µm. (C) PIH1D1-GFP at different time points of invadosome formation in NIH3T3-Src LifeAct-mCherry cells by video-microscopy. Scale bar, 15 µm (n = 3; t test; p < 0.01; p < 0.001; **p < 0.0001).

**Figure S3.**
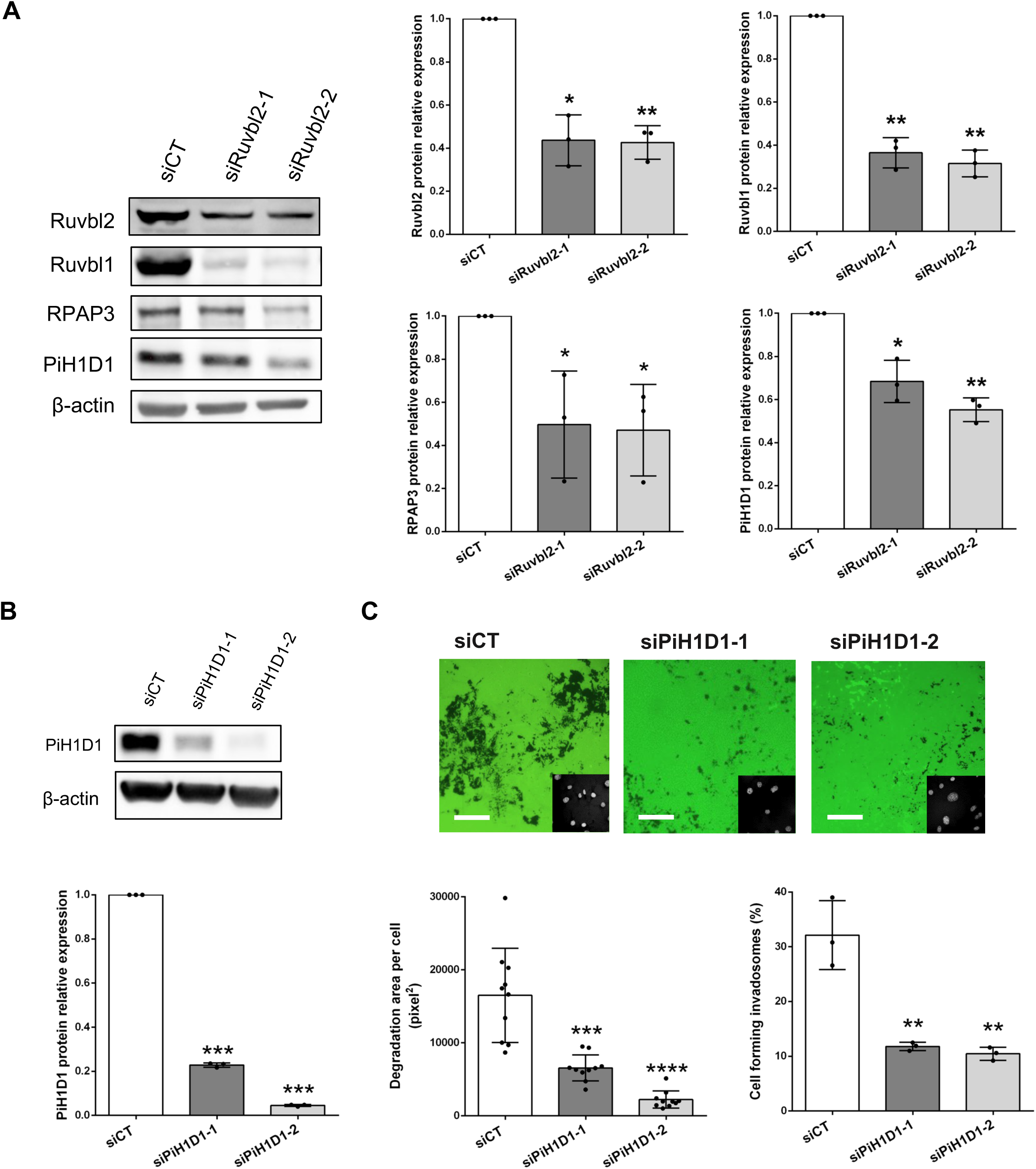
**The R2TP complex is coordinately regulated and PIH1D1 is required for invadosome formation and activity**, related to Figure 2. (A) Western blot analysis of RUVBL2, RUVBL1, RPAP3 and PIH1D1 in RUVBL2-depleted NIH3T3-Src cells. Quantification, right. (B) Western blot analysis of PIH1D1 in PIH1D1-depleted NIH3T3-Src cells. Quantification below. (C) In situ zymography of these cells, with quantification of degradation and invadosome formation below. Scale bar, 150 µm (n = 3; t test; *p < 0.05; **p < 0.01).

**Figure S4.**
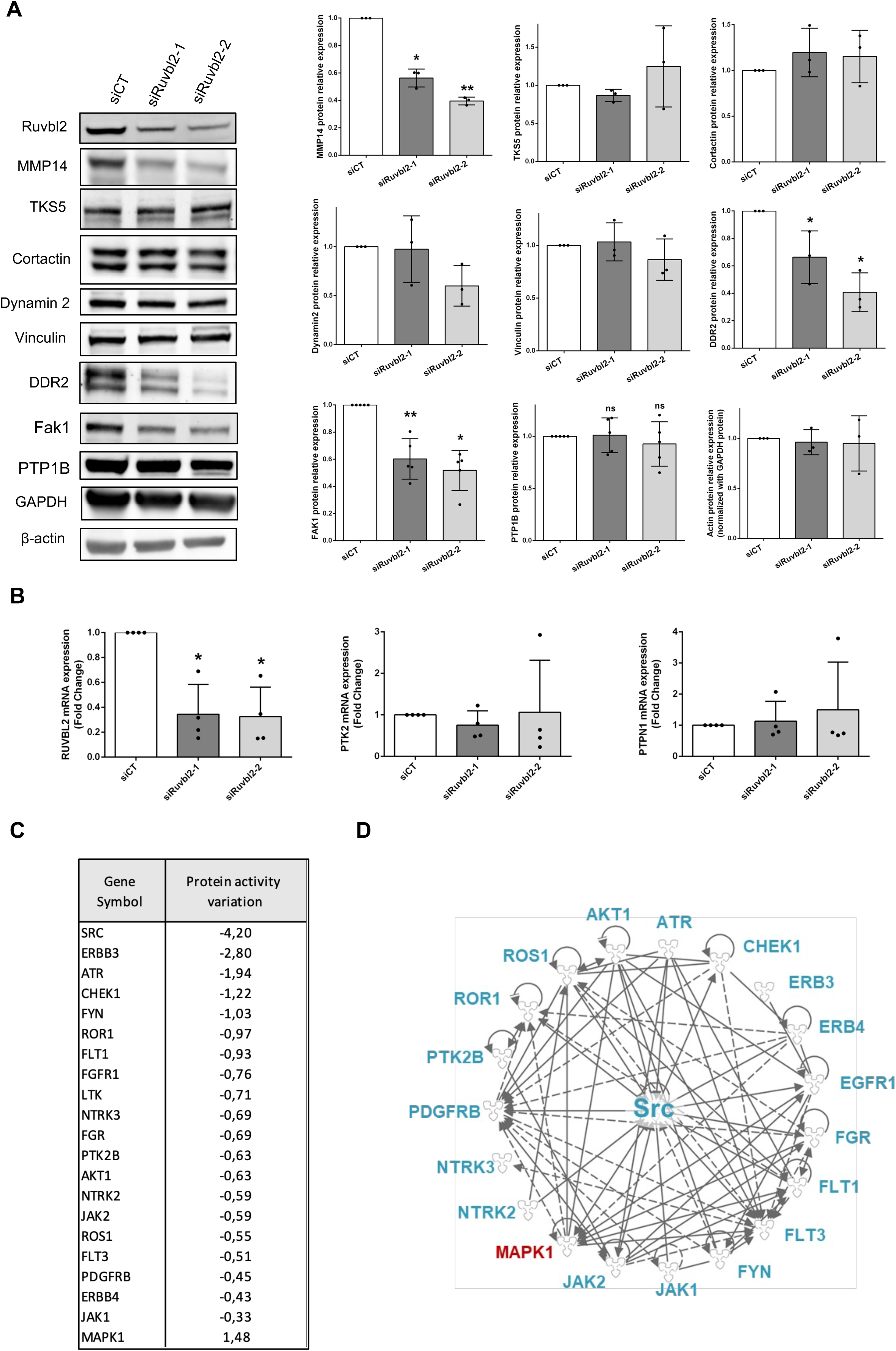
**RUVBL2 depletion affects invadosome proteins and kinase activity**, related to Figure 3. (A) Western blot analysis of MMP14, TKS5, cortactin, dynamin 2, vinculin, DDR2, FAK1, PTP1B and GAPDH in RUVBL2-depleted NIH3T3-Src cells. Quantification, right. (B) RT-qPCR analysis of Ptk2 and Ptpn1 transcripts under the same conditions. (C) Kinases whose activity is modified upon RUVBL2 depletion, from a PamGene PTK assay, shown with gene names and activity variation. (D) Network representation showing that these kinases belong to the Src environment and can interact with it directly or indirectly, generated with Ingenuity Pathway Analysis (n = 3; t test; p < 0.05; p < 0.01; p < 0.001).

**Figure S5.**
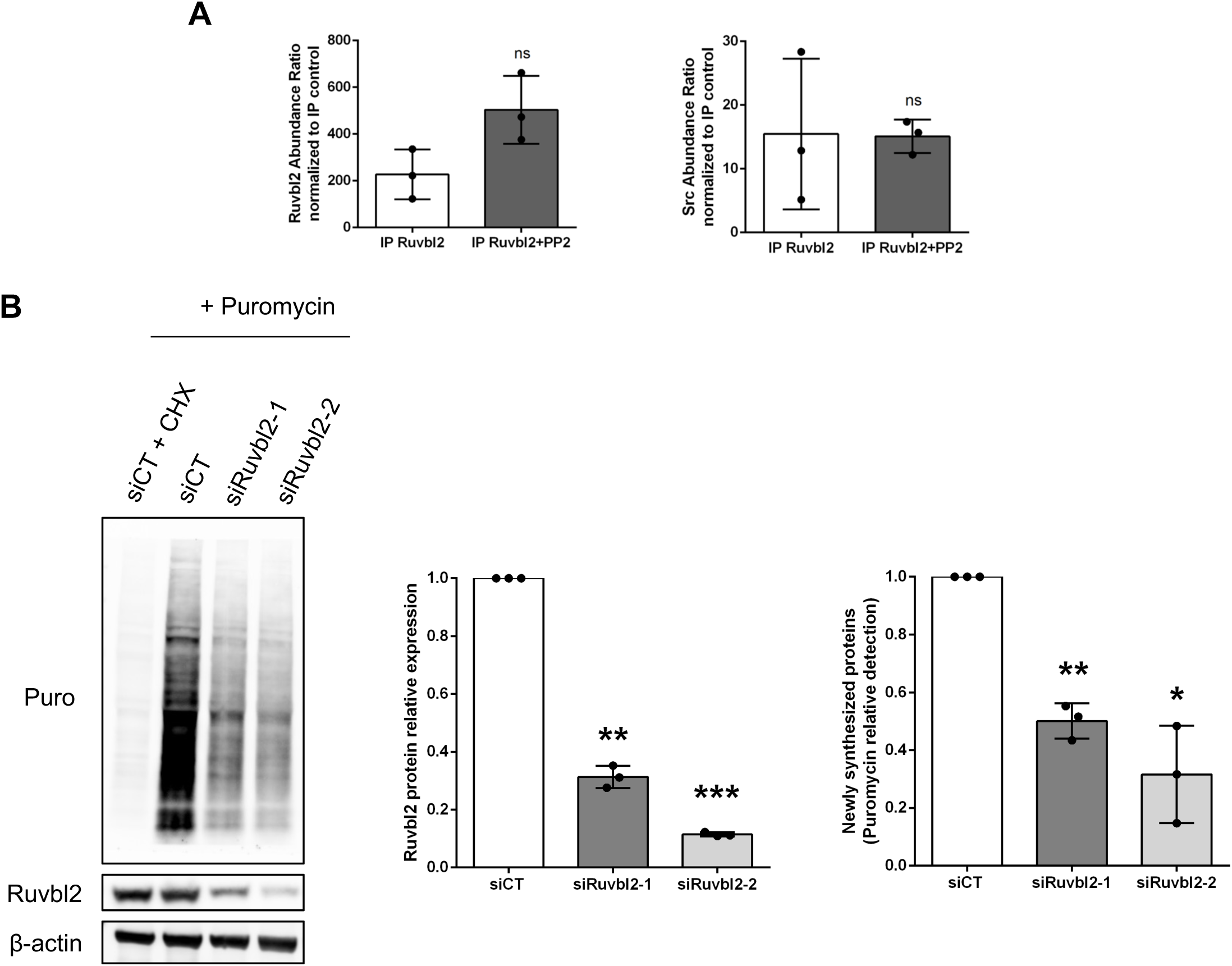
**Src-dependent RUVBL2 partners are involved in protein synthesis**, related to Figure 3. (A) Western blot analysis of RUVBL2 in lysates after Flag immunoprecipitation in NIH3T3-Src cells transfected with CMV-Flag or RUVBL2-Flag and treated or not with PP2. (B) Western blot analysis of RUVBL2 and puromycin incorporation, corresponding to newly synthesised proteins in the SUnSET assay. Cycloheximide (CHX) was used as a negative control. Quantification, right (n = 3; t test; p < 0.05; p < 0.01; p < 0.001).

**Figure S6.**
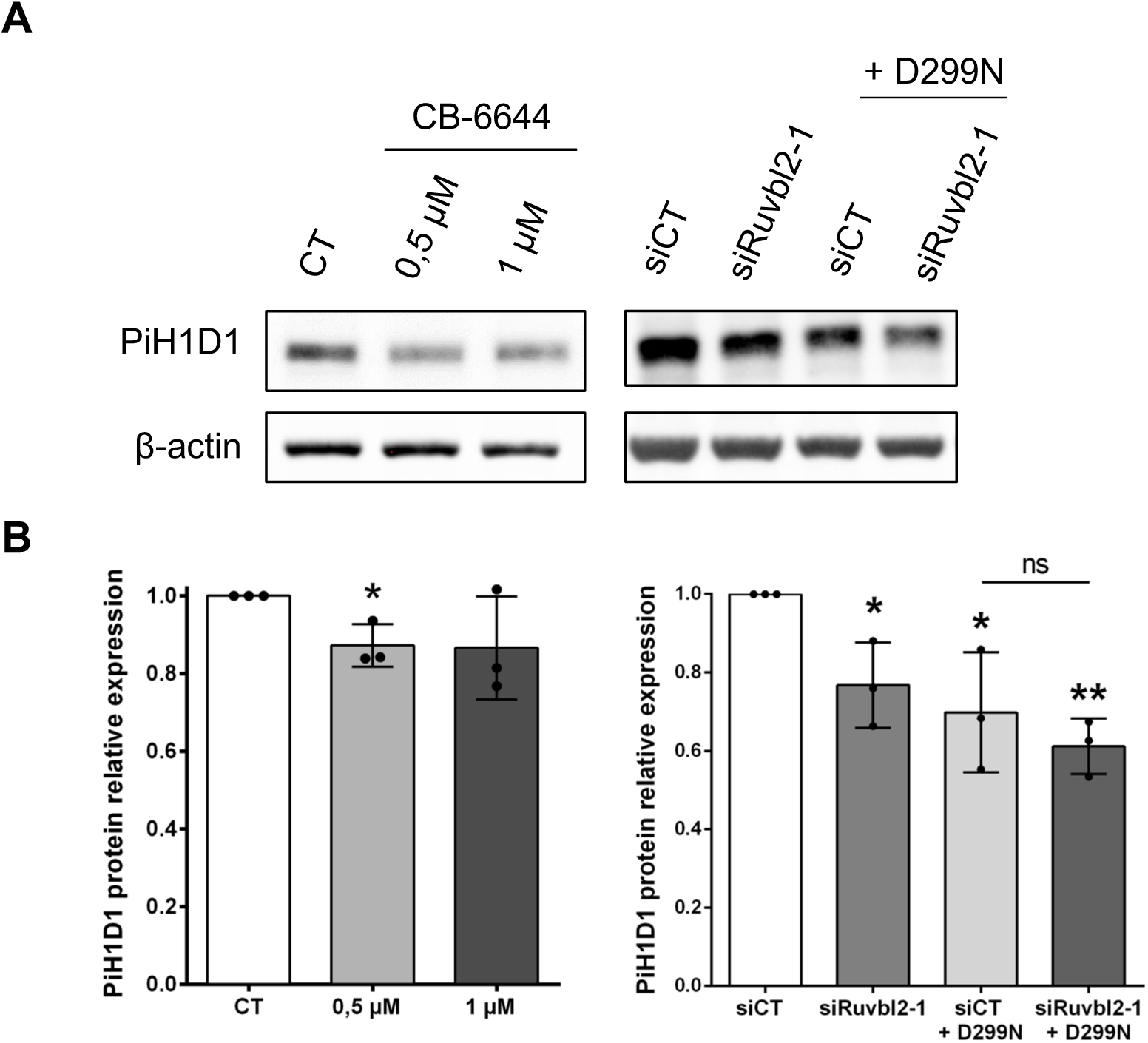
**Inhibition of RUVBL2 ATPase activity decreases PIH1D1 protein levels**, related to Figure 5. (A) Western blot analysis of PIH1D1 in NIH3T3-Src cells treated with CB-6644 (0.5 and 1 µM) or co-transfected with the RUVBL2 D299N mutant. (B) Associated quantification. Video S1. RUVBL2 dynamics in rosette invadosomes, related to Figure 1. Seventy-nine-minute time-lapse of living NIH3T3-Src cells expressing LifeAct-mCherry (red) and RUVBL2-GFP (green).

**Table S1**. Proteins significantly deregulated upon RUVBL2 depletion, related to Figure 3.

**Table S2**. RUVBL2 interactors altered by Src inhibition, related to Figure 3.

---

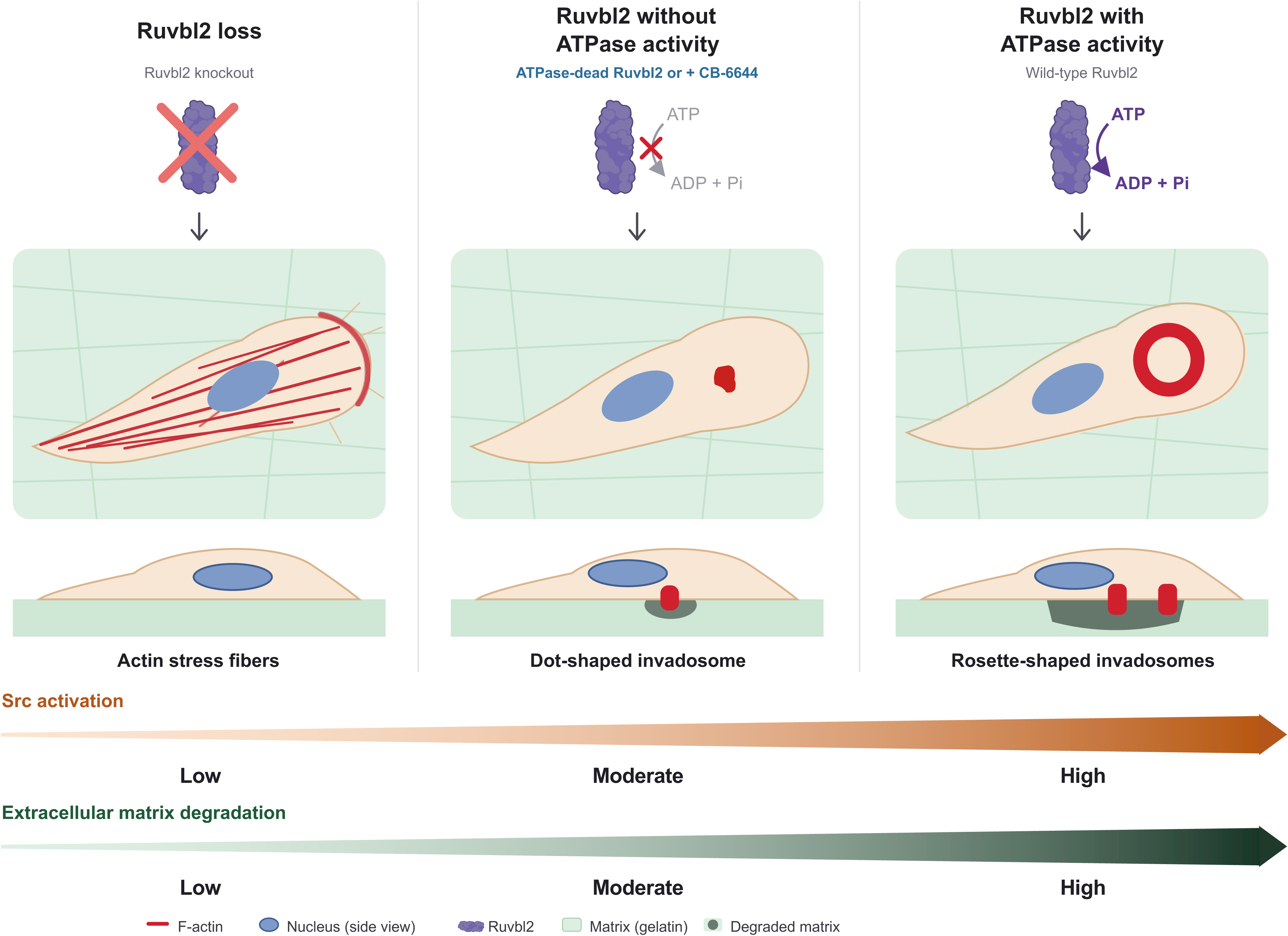

